# Screening for inborn errors of metabolism using untargeted metabolomics and out-of-batch controls

**DOI:** 10.1101/2020.04.14.040469

**Authors:** Michiel Bongaerts, Ramon Bonte, Serwet Demirdas, Ed H. Jacobs, E. Oussoren, Ans T. van der Ploeg, Margreet A.E.M. Wagenmakers, Robert M.W. Hofstra, Henk J. Blom, Marcel J.T. Reinders, George J. G. Ruijter

## Abstract

**Motivation:** Untargeted metabolomics is an emerging technology in the laboratory diagnosis of inborn errors of metabolism (IEM). In order to judge if metabolite levels are abnormal, analysis of a large number of reference samples is crucial to correct for variations in metabolite concentrations resulting from factors such as diet, age and gender. However, a large number of controls requires the use of out-of-batch controls, which is hampered by the semi-quantitative nature of untargeted metabolomics data, i.e. technical variations between batches. Methods to merge and accurately normalize data from multiple batches are urgently needed.

**Methods & results:** Based on six metrics, we compared existing normalization methods on their ability to reduce batch effects from eight independently processed batches. Many of those showed marginal performances, which motivated us to develop *Metchalizer*, a normalization method which uses 17 stable isotope-labeled internal standards and a mixed effect model. In addition, we propose a regression model with age- and sex as covariates fitted on control samples obtained from all eight batches. *Metchalizer* applied on log-transformed data showed the most promising performance on batch effect removal as well as in the detection of 178 known biomarkers across 45 IEM patient samples and performed at least similar to an approach using 15 within-batch controls. Furthermore, our regression model indicates that 10-24% of the considered features showed significant age-dependent variations.

**Conclusions:** Our comprehensive comparison of normalization methods showed that our *Log-Metchalizer* approach enables the use out-of-batch controls to establish clinically-relevant reference values for metabolite concentrations. These findings opens possibilities to use large scale out-of-batch control samples in a clinical setting, increasing throughput and detection accuracy.

**Availability:** *Metchalizer* is available at https://github.com/mbongaerts/Metchalizer/

## Introduction

Screening of patients suspected for inborn errors of metabolism (IEM) is currently based on measuring panels of specific groups of metabolites like amino acids or organic acids using a number of different tests and techniques such as ion-exchange chromatography, LC-MS/MS and GS-MS. This targeted approach with several different tests is time consuming and limited in the number of metabolites being analyzed. Untargeted metabolomics using High Resolution Accurate Mass Liquid Chromatography Mass Spectrometry (HRAM LC-MS) can detect hundreds to thousands of metabolites within one test, and, as a consequence, receives increasing interest to be used in IEM screening (Miller, et al., 2015) (Coene, et al., 2018) (Körver-Keularts, et al., 2018) (Haijes, et al., 2019) (Bonte, et al., 2019). Moreover, untargeted metabolomics can also reveal new biomarkers or increase our understanding of disease mechanism when exploited in epidemiological studies (Glinton, et al., 2019).

In traditional targeted diagnostic laboratory tests hundreds of reference samples are required to establish robust reference intervals. When using untargeted metabolomics the establishment of reference values is complicated due to the semi-quantitative nature of the data owing to several sources of variation like injection volume, retention time, temperature, or ionization efficiency in the mass spectrometer that cannot easily be amended. Moreover, these variations are even larger between different measurement runs in which a batch of samples is being measured simultaneously, hampering the resemblance between different batches. As a result, within-batch variation is smaller than between-batch variation. Therefore, to conquer these batch effects, current approaches include reference samples in each single batch of measurements (Miller, et al., 2015) (Coene, et al., 2018) (Haijes, et al., 2019) (Körver-Keularts, et al., 2018) (Bonte, et al., 2019) to improve detection sensitivity (due to tighter reference values as a result of lower variation in the in-batch reference samples).

Clearly, this reduces the throughput efficiency of IEM screening as the number of patient samples that can be included in a batch is considerably lower when the reference samples need to be measured as well. But, more importantly, the number of reference samples in one batch might fall short in the establishment of adequate reference ranges as variations in certain metabolites are not captured well enough in the relatively small reference panel. For example, factors like age, sex and BMI can affect abundancies of metabolites, and, to establish reliable reference ranges, one thus needs to correct for these factors by using a large number of reference samples (Chaleckis, et al., 2016) (Rist, et al., 2017) (Yu, et al., 2012). Consequently, for reliable untargeted metabolomics in clinical testing, a large set of reference samples is needed, while for throughput efficiency a small set is preferred. Altogether, this calls for an approach that can establish reference values based on reference samples being measured in several batches (out-of-batch controls).

When relying on reference samples from different batches, one needs to correct for the batch effects to obtain reliable estimates for the reference ranges. This is generally solved by normalization methods and some have already been proposed within the context of untargeted metabolomics and mass spectrometry (Veselkov, et al., 2011) (Li, et al., 2017) (Välikangas, et al., 2016). Only a few groups have used out-of-batch controls to determine the reference values and used relatively simple normalization techniques like median scaling (Miller, et al., 2015), using a reference internal standard per metabolite (Körver-Keularts, et al., 2018) or using anchor samples (Glinton, et al., 2019). However, there has not been an extensive exploration of normalization techniques within the context of diagnostic testing for IEM’s.

We explore several known normalization methods on their ability to remove batch effects and to detect biomarkers from patients with known IEM. Furthermore, we introduce a new normalization method, which we called *Metchalizer*, which uses internal standards and a mixed effect model to remove batch effects. As this allows for a large set of (out-of-batch) reference samples, we also explore a regression model that uses age and sex as covariates to correct for potential age and sex effects on the reference values. Using the regression model combined with the *Metchalizer* normalization, we achieve similar performances in biomarker detection compared to the use of within-batch controls. Hence, this opens the possibility to increase the throughput of untargeted metabolomics in IEM screening as well as including more complex confounder strategies.

## Materials and methods

### Untargeted metabolomics datasets

Human plasma samples of 260 control samples and 53 IEM patients were measured over eight batches over the period 10-12-2018 to 03-05-2019 (Bonte, et al., 2019) having in total 33 unique IEMs. For every patient a technical triplicate was included. A QC (Quality Control) sample was included in all eight batches and more than four technical replicates were present in every batch. Since the QC sample was a commercial sample, the sample differed in concentration of several metabolites when compared to the (average) concentrations of the human plasma samples analyzed in these datasets. Features were annotated as described in Bonte et al. (Bonte, et al., 2019). Note that within each batch about 30 normal controls have been measured, which allows us to establish reference values based on within-batch controls, whereas the controls being measured for the other (seven) batches can be used for out-of-batch strategies. In this study we will refer to ‘feature’ as being either a single m/z-value (with unique retention time) or a merge of multiple features, where the adduct type and/or isotope was determined with corresponding neutral mass and consequently merged to a single feature.

The following internal standards have been added to each batch to facilitate normalization based on these internal standards: 1,3-^15^N uracil (+/−), 5-bromotryptophan (+/−), D_10_-isoleucine (+/−), D_3_-carnitine (+/−), D_4_-tyrosine (+/−), D_5_-phenylalanine (+/−), D_6_-ornithine (+), dimethyl-3,3-glutaric acid (+/−), ^13^C-thymidine (+/−), D_4_-glycochenodeoxycholic acid (−), where + indicates positive ion mode, and – indicates the negative ion mode.

### Data processing

Previous pre-processing steps (alignment, peak picking etc.) were performed per batch using Progenesis QI v2.4 (Newcastle-upon-Tyne, UK) (Bonte, et al., 2019). In-house software was developed to match features from each batch to a reference batch which in this case was the fifth batch when sorting on chronologically order. Chromatograms between batches were initially aligned to the reference batch by using lowess regression where features were matched based on retention time difference, m/z-value and median abundancy difference similar to the criteria described below.

Matching features was performed based on several criteria:

1. When features were annotated in reference batch and the batch being merged, these features were pooled to the merged dataset.
2. When MS/MS spectra were present for a potential matching pair of features, the cosine similarity metric was calculated and had to be > 0.8.
3. Retention time difference in percentage was calculated between potential matches, and had to be < 2.5%.
4. Progenesis QI determined per feature an isotope distribution and we required sufficient overlap of these distributions between potential matching pairs. This was determined by calculating a difference in percentage between each bin of this distribution. The maximum difference of these bins had to be < 50%.
5. As we expect matching features to have similar within-batch median abundancies (despite of batch effects), we calculated the differences between these medians in percentages, which had to be < 300%.
6. Neutral masses were known for the matching pair but not the MS/MS spectra, the ppm-error had to be < 1.
7. m/z-values were known for the matching pair but not the MS/MS spectra and neutral masses, the ppm-error of between the m/z-values had to be < 1.

Features matching multiple other features in the reference batch were discarded (and vice versa). The resulting merged dataset contained only features which were matched across all eight batches.

### Quantitative evaluation set

For the evaluation of the normalization methods, the following 16 metabolites were quantitatively (μmol/L) measured in two separate assays: leucine (+/−), C0 | L-carnitine (+/−), methionine (+/−), C2 | acetylcarnitine (+), 5-aminolevulinic acid/4-hydroxyproline (+), serine (+/−), citrulline (+/−), aspartic acid (+), glutamine (+/−), (allo)isoleucine (+/−), proline (+/−), tyrosine (+), phenylalanine (+/−), taurine (+/−), asparagine (+/−), arginine (+/−). Amino acids were determined by ion-exchange chromatography according to protocols described by the manufacturer (Biochrom). Free carnitine and acylcarnitines analysis was performed as described by Vreken et al. (Vreken, et al., 2002).

### Normalization methods

#### Initial transformations

Prior to normalization raw abundancies were for some methods transformed using a log-transform or Box-Cox transformation. The latter was given by:

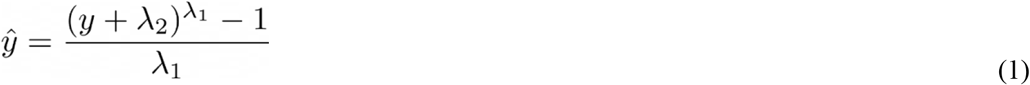

with λ_1_ = 0.5 and λ_2_ = 1. If an initial transformation was applied this was indicated in the name of the (normalization) method, where ‘BC-’ refers to the Box-Cox transformation and ‘Log-’ to the log transformation. When no transformation was performed this was indicated with ‘None-’.

### Normalization by Metchalizer

*Metchalizer* assumes a linear mixed effect relationship between the abundancies of the internal standards and the feature of interest. Since the internal standards were expected to be correlated, we represented them by an orthogonal set of covariates. These covariates are obtained as the Latent Variables (LV) from the Partial Least Squares (PLS) of the set of internal standard abundancies (represented in matrix **X**) and the (categorical) information about which sample belonged to which batch (represented by matrix **Y**). The number of LV’s were chosen from the metric *I(K)*:

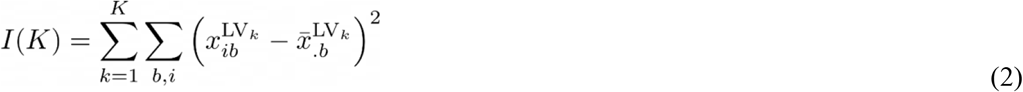

where 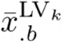 is the center of batch *b* in the direction of LV_*k*_. We selected that *K* for which *I(K)* reached 75% of its maximum value.

The mixed effect model then considers the LV’s as fixed effects and all variations not explained by the LV’s is considered as (random) batch effects:

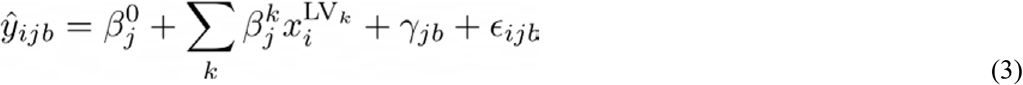

with 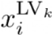 indicating the covariate (score) of the *k*^th^ Latent Variable (LV) of sample *i*. *γ*_*jb*_ is the (random) batch intercept for feature *j*. Note, that when the LV’s are sufficient in explaining *y*_*ijb*_ the random intercept γ_jb_ will not contribute much. Before fitting the model, we remove outlier samples per batch *b* and feature *j* based on their within-batch Z-score (|Z| > 2) determined from all samples in that batch. These Z-scores were different than the Z-scores defined in other parts of this study.

The batch corrected abundancy then becomes:

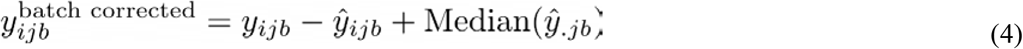

### Normalization by Best Correlated Internal Standard

The internal standard, *m*, that best correlates with a feature *j* is being used to normalize the abundances of feature *j*. The correlation is measured within each batch using the spearman correlation between feature *j* and each internal standard individually across all samples and subsequently averaged across all eight batches. The internal standard which (positively) correlated the best was used for normalization according:

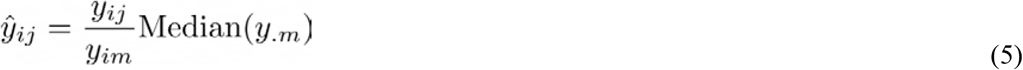

with *m* being the best correlated internal standard.

### Normalization methods from literature

We compared *Metchalizer* with a number of different normalization methods. For a description we refer to the original articles, here we only specify our settings:

#### Anchor (Glinton, et al., 2019)

*Anchor* assumes a linear response between the features in the anchor samples and samples in the batch. An anchor sample is a fixed sample which is analyzed in all eight batches, and was included more than four times in each batch. Normalization was performed per batch by dividing each feature by the median of the anchor samples for that same feature per batch [1]. In this study we used our QC samples as the anchor samples.

#### CRMN (Redestig, et al., 2009)

We used function normFit from the *crmn* R package with input argument “`crmn`” and `ncomp=3`. As a design matrix we chose QC samples versus human plasma’s.

#### EigenMS (Karpievitch, et al., 2015)

QC samples and plasma samples were treated as two different groups. We chose three ‘eigentrends’.

#### Fast Cyclic Loess (Ballman, et al., 2004)

We used the *normalizeCyclicLoess* function from the *limma* R package using the method `“fast”` and `span=0.7`.

#### NOMIS (Sysi-Aho, et al., 2007)

We used the function *normFit* from the *crmn* R package with input argument `“nomis”`.

#### PQN (Filzmoser & Walczak, 2014)

*PQN* was implemented as described by Filzmose et al. The reference spectrum was given by the median of every feature *j*.

#### RUV (Livera, et al., 2015)

We used the function *RUVRand* with `k=8` from the *MetNorm* R package.

#### VSN (Huber, et al., 2002)

We used the *vsn* R package using the *vsn2* function.

### Evaluation of normalization methods

Six metrics were used to evaluate the performance of normalization methods.

#### WTR_j_ score

The WTR score (**W**ithin variance **T**otal variance **R**atio) calculates the ratio between the ‘overall’ within-batch variance and the total variance from the QC samples:

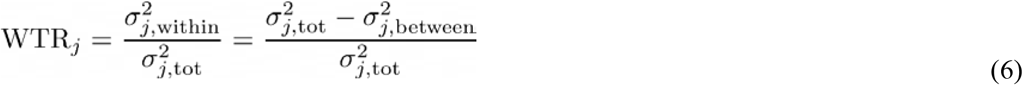

where *σ*_*j*_ is the variance of all eight batch averages for metabolite *j* in the QC samples, and 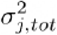 the ‘overall’ variance based on all QC samples. The WTR score is between 0 and 1. As we would like batch averages to be similar for the QC samples (resulting in *σ*_*j*_, between approaching zero), we are interested in WTR scores close to one.

#### Δ*R* score

Since normalization might also lead to the removal of variations of interest (for example biological variations), we tested whether the ranks of the features ordered by their abundancies within the QC samples were preserved after normalization. Per feature *j*, we determined the average rank the feature is assigned across all QC samples (across all batches) for both the raw abundancies 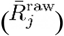 as well as the normalized abundancies 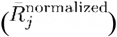. The Δ*R*_*j*_ score then looks at the difference in rank positions due to normalization per feature *j*:

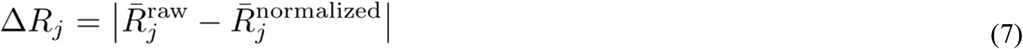

Δ*R*_j_ ∈ [0, *p*], with *p* the number of features. Lower Δ*R*_*j*_ values indicate a better preservation of the ranks of the normalization method.

#### Spearman score

For the set of 16 quantitatively measured metabolites, we calculated the Spearman correlation between their quantitative measurements and the normalized abundancies. Overall normalization performance could be judged based on the median Spearman score of these 16 scores, having scores ∈ [−1, 1]. Higher values indicate better resemblance with the quantitative measurements.

#### R^2^ score

The R^2^ between the quantitative measurements and the normalized abundancies of the 16 quantitatively measured metabolites. Overall performance could be judged from the median R^2^ score, with scores ∈ [0, 1]. Higher values indicate better (linear) fits with the quantitative measurements.

#### QC prediction score

Since the QC samples were different from the human plasma samples in terms of concentrations for several metabolites/features, we expect this difference to be observed in the first few principal components (PCs) of a Principal Component Analysis (PCA) analysis applied to all features (excl. standards). We fitted a logistic function using the first four PC’s as covariates and with class labels: ‘human plasma’ and ‘QC’. The fitted model returns per sample a probability of belonging either to the class ‘human plasma’ or ‘QC’. The probabilities for all samples are averaged into the *QC prediction score* ∈ [0, 1] Increasing normalization performances should result in higher scores, as QC - and human plasma samples should be nicely separated. We used *LogisticRegression* from the Python package *scikitlearn* with parameters `penalty=‘l1’`, `solver=‘saga’`, `multi_class=‘auto’`, `max_iter=10000` (Pedregosa, et al., 2011).

#### Batch prediction score

Increasing normalization performances should result in less batch clustering when examining the first few PC’s of the PCA analysis (see *QC prediction score*). We fitted a logistic function for each batch versus all other seven batches using the first four PC’s as covariates and obtained the probability scores for all human plasma’s having the correct batch label. These scores were than averaged for all human plasma samples into a *batch prediction scores* ∈ [0, 1]. Scores closer to 1 indicate decreased normalization performances since batch separation is (still) present.

### Methods to determine aberrated metabolic abundancies

Reference values for metabolites were determined by using a Z-score methodology: a set of reference values was Z-transformed (corrected for mean and divided by the standard deviation) which was then assumed to be normally distributed. Aberrations can then be called by considering significant Z-scores using a chosen cutoff level. We use four different methods to determine the Z-scores.

#### Method *15in*: best matching controls within batch

Z-scores were calculated by selecting 15 control samples originating from the same batch as the patient based on age and sex as described in Bonte et al. (Bonte, et al., 2019).

#### Method *15out*: best matching controls from other batches

Z-scores were calculated similarly as in *method 15in* using explicitly 15 out-of-batch controls. Note, that since there a more out-of-batch controls than within-batch controls that age and sex matching can be done more accurately.

#### Method *All controls*

This method used all available control samples from all eight batches, including within-batch controls, for Z-score calculation.

#### Method *Regression*

We fitted a linear model on all 260 available controls excluding outliers which were first removed based on their within-batch |Z-score| > 3, this Z-score is different from other Z-scores mentioned in this study, and only used to remove outliers. The regression model is given by:

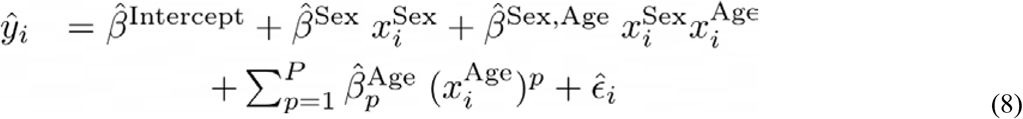

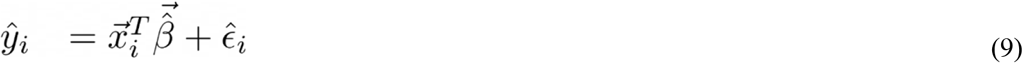

where 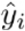 is the predicted (normalized) abundancy of feature *j* for sample *I*, 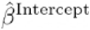 is an intercept. 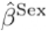, 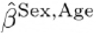 (interaction) and 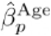 indicate slopes. *P* is the degree of the polynomial used for regression on age and set to *P*=3 in this study. 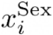 is 1 for women and 0 for men. 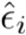 is the estimated error. The latter expression is the model in vector notation with 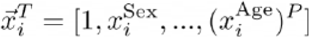.

The coefficients were determined from the OLS estimator:

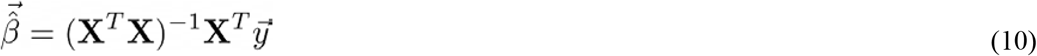

where the rows of **X** are given by 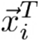 and the variance in 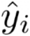 is determined by the variance in 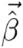 and the variance in 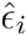:

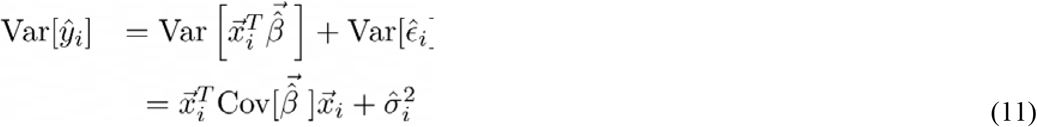

The covariance matrix of 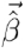 is given by:

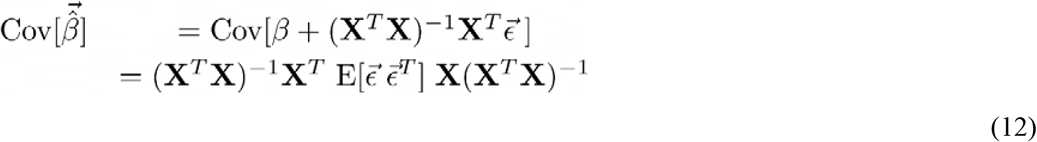

with 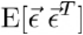 estimated according:

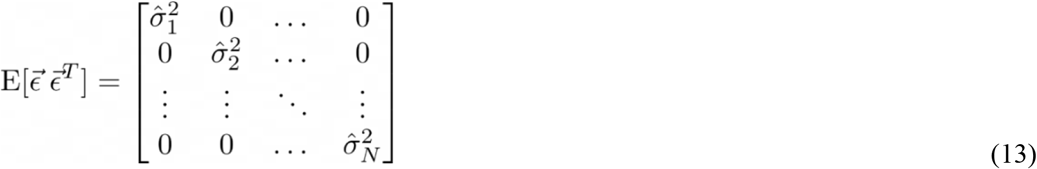

Since we expected 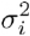 to be dependent on age (neglecting sex), we do estimate 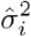 differently from a weighted mean on the squared residuals:

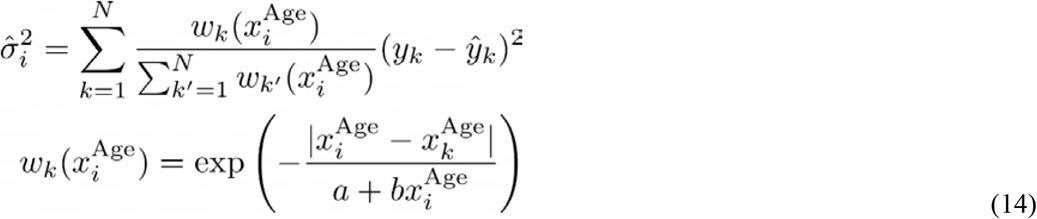

where *a* and *b* determine how the weights decay (*a*) or increase (*b*) over age (we set *a, b = 1* years). Z-scores were obtained by subtracting the predicted average 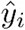 and dividing by the variance 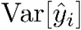 (Equation 11).

#### Significance of regression coefficients

Significance of the regression coefficients (Equation 8, 9) was obtained by considering the statistic:

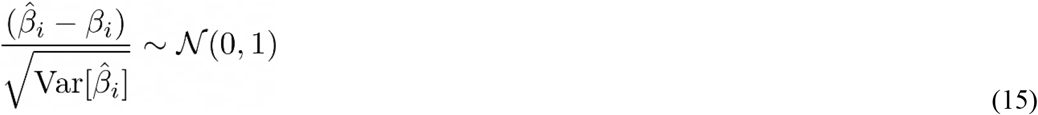

The variances of the coefficients were found in the diagonal elements of 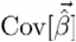 (Equation 13). We tested the hypotheses that *β*_*i*_=0 with a two-tailed test. A robust p-value was obtained from a bootstrap procedure by taking the median p-value from a series of p-values obtained from 50 bootstraps on the above test statistics taking 95 % of the data each bootstrap.

### Final Z-scores

Since the patient samples were measured in triplicate, we determined the final Z-scores from the average of these three Z-scores (Bonte, et al., 2019). These average Z-score were determined for all Z-score methods i.e. *15in, 15out, All controls, Regression* and IEM patient.

### P-values from Welch’s t-test

As an alternative to using the (average) Z-scores we also considered the p-values obtained from the Welch’s t-test to be informative, as it indicates whether the mean of triplicates differs significantly from the population average. Note that the triplicate was expected to have only technical variance whereas the reference population has variance consisting of technical-plus biological variance. For every Z-score method (*15in, 15out, All controls, Regression*) these p-values were obtained per feature (and patient).

When using the regression model, we used an adjusted Welch’s t-test assuming that variance in the estimate of the average of the population (which is Z=0) was negligible:

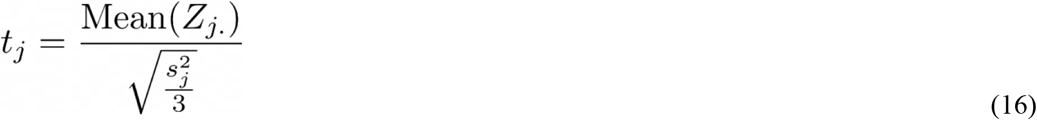

where *s*_*j*_ is the sample standard deviation of the triplicate Z-scores, Mean(*Z*_*j*_) indicates the average of the triplicate for feature *j*.

### Detection of the expected IEM biomarkers

To explore how normalization and the method of determining these Z-scores (*15in, 15out, All controls* and *Regression*) affected the detection of biomarkers, we plotted the number of detected biomarker of the known IEM patients against the average number of detected features per patients for various (final) Z-score and p-value cutoff levels, similar to a ROC curve. Improved biomarker detection was believed to increase the area under the ROC(-like) curve (AUC).

Establishing this ROC curve was done by assigning a status for every biomarker (if present and annotated in the MS-data). A database was established containing the expected biomarkers for each IEM including the expected Z-score sign (up or down regulated) as can be found in supplement S5 Table 5. For every IEM patient, we assigned for all expected biomarkers the status ‘positive’ or ‘negative’. The status ‘positive’ was assigned when 1) |Z-score| > Z_abnormal_, and 2) the sign of the Z-score corresponded with the expected sign for that biomarker in the IEM patient. Criteria 1 and 2 were also used for the ROC-curve created by the p-values. When a biomarker was found in both positive and negative ion mode, the Z-score(s) from the mode having the largest population average abundancy was taken. The average number of detected features (per patient) was obtained by considering features from both ion modes.

Some of the expected biomarkers were not matched across all eight batches and therefore were absent in the merged dataset and analysis in this study. In the merged dataset, we obtained 178 patient-biomarker combinations (one patient could have multiple biomarkers) associated with 45 patients (hence, for 8 IEM patients no biomarkers were found in the merged dataset).

## Results

### Batch characteristics

Eight untargeted metabolomics runs/batches were merged containing 260 control samples and 53 IEM patients, together having 33 unique IEMs. After merging, 773 positively ionized features were obtained, among which 121 were annotated, and 598 negatively ionized features were attained with 106 annotated features. We only included features which were merged across all eight batches to ensure consistency among the findings. Intra-batch coefficients of variation (CV) on 17 (internal and external) standards were smaller (median CV=14%) than inter-batch CV’s (median CV=27%) indicating that batch effects were present (for more details see S1). Principle Component Analysis (PCA) further elucidated the presence of batch effects as shown in Figure 2A, showing the first three PC’s for the log-transformed raw abundancies (*Log-Raw*).

### Comparing normalization methods

We investigated the performance of several normalization methods on batch effect removal by evaluating multiple metrics based on quantitative measurements, the Quality Control (QC) samples and PCA analysis (see Methods and Figure 1). Some normalization methods were excluded from the following analysis because of their marginal performance on the considered metrics (as evaluated in supplement S2).

**Figure 1.**
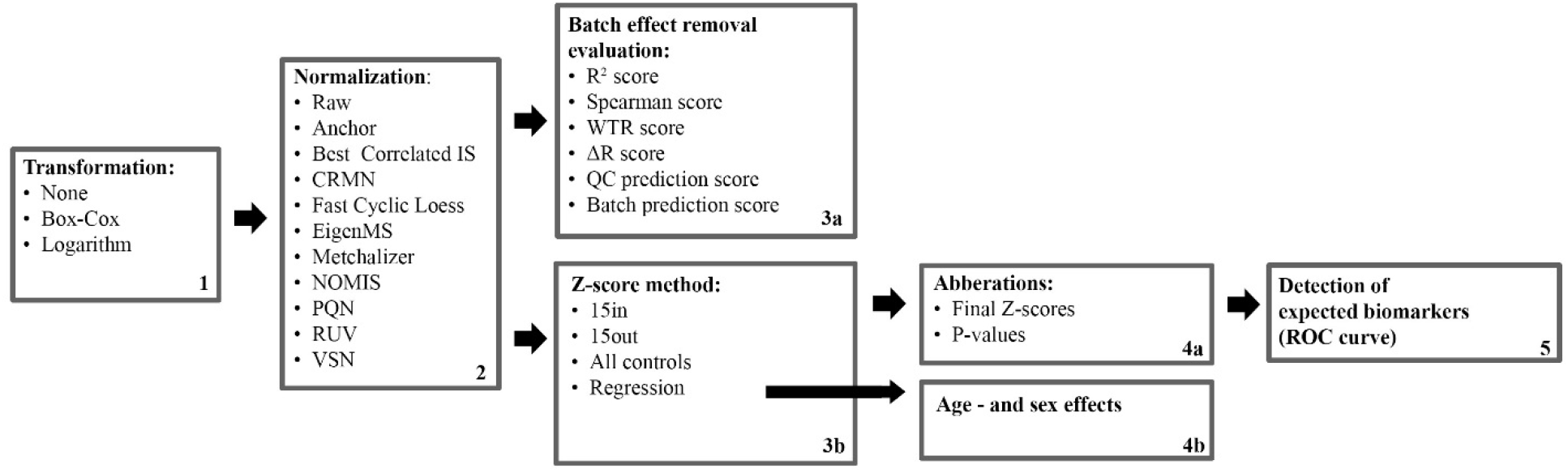
Flow diagram of different methods used in this study. 1) An initial transformation was applied. 2) A normalization method was applied. 3a) Multiple metrics were calculated to investigate batch effect removal. 3b) Normalized data was used to determine Z-scores for IEM patients using different (control) reference methods. 4a) Final Z-scores were calculated together with p-values. 4b) Regression analysis on all features/biomarkers was used to explore age-and sex dependency of abundancies. 5) Detection of the expected biomarkers was investigated using a ROC-like curve for Z-scores and p-values

**Figure 2.**
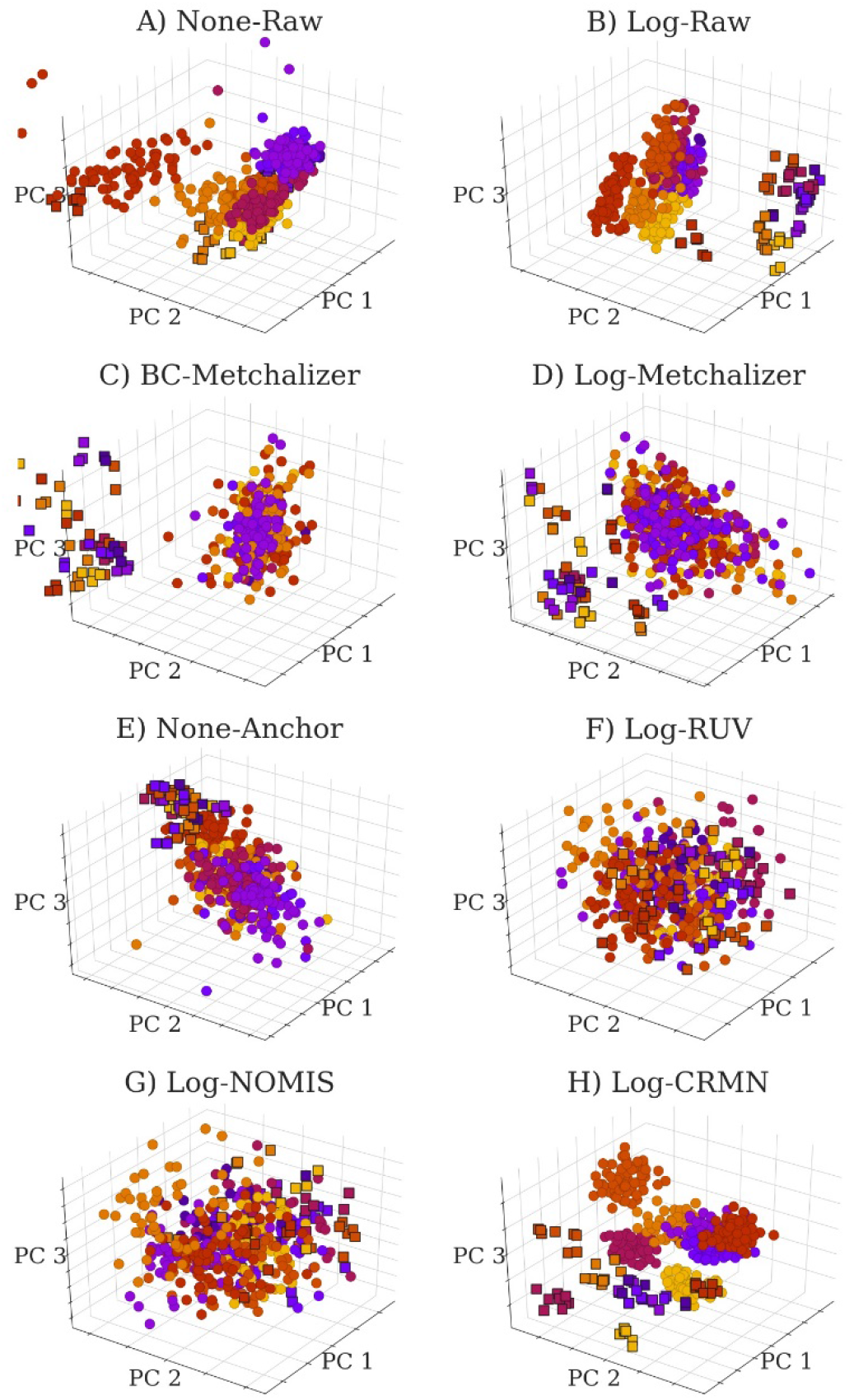
PCA plots for raw data and normalized data as indicated by the title of each panel. Each batch is indicated with a unique color. PCA was performed on 758 features (excluding the internal – and external standards) in positive ion mode. The squares indicate QC samples whereas the circles indicate patients and controls samples.

#### Reduced batch effects

We visually observe in the PCA plots that most normalization methods reduced batch effects since batch clustering seemed to be reduced after normalization (Figure 2), and is confirmed when looking at the *batch prediction score* (Figure 3A) showing lower scores for normalized abundancies when compared with the raw data (*None-Raw* or *Log-Raw*). *BC-Metchalizer*, *Log-Metchalizer* and *None-Anchor* had the lowest *batch prediction scores*, with a median score of 0.13 (0.13), 0.14 (0.14), 0.17 (0.16) for positive (negative) ion mode respectively.

**Figure 3.**
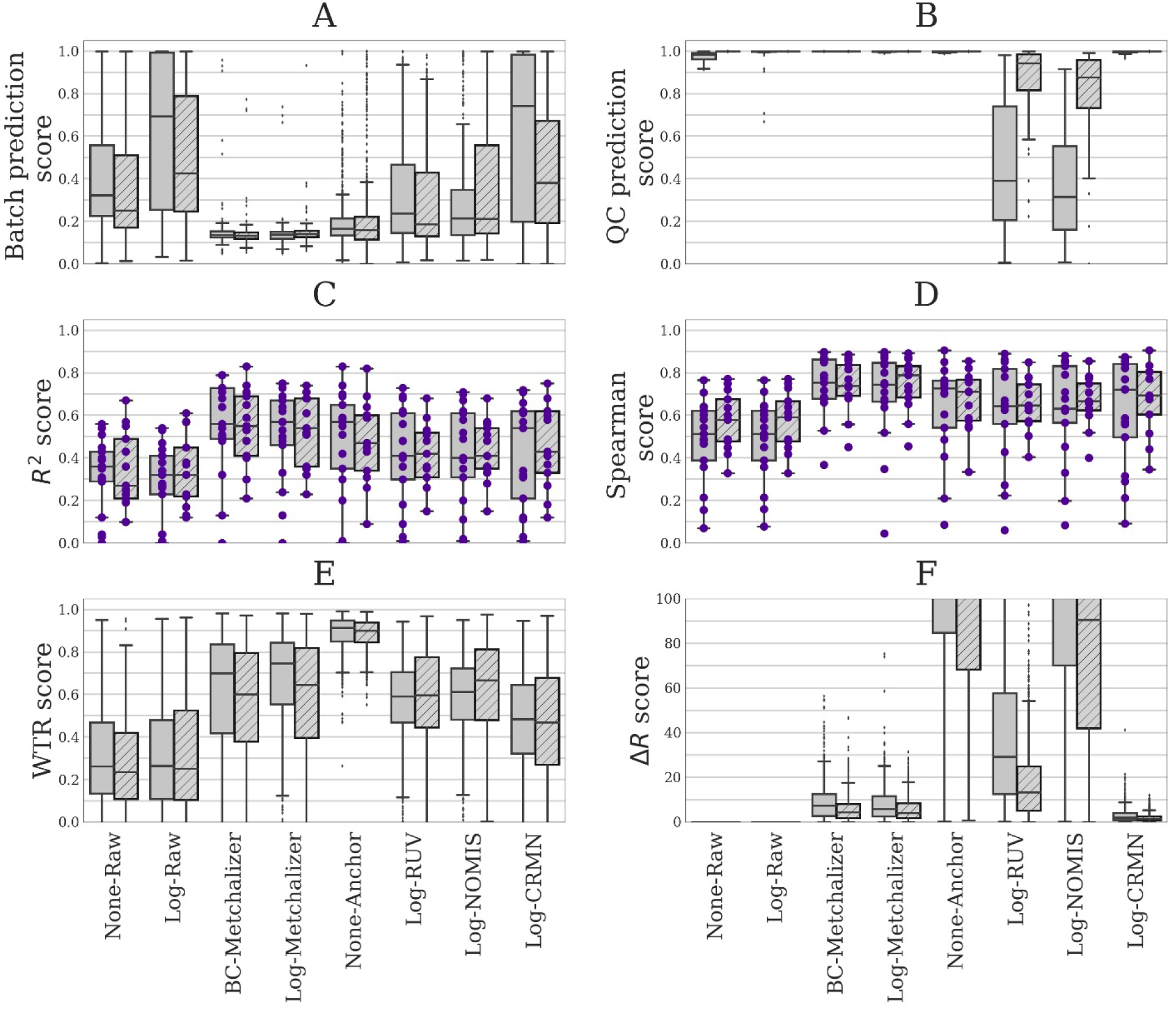
Six different performance metrics for batch effect removal (see Methods for more details). Data from positive – or negative ion mode is indicated by plain and stripped boxplots, respectively. A) *Batch prediction score* measures the presence of batch effects in the first four PC’s from PCA analysis. B) *QC prediction score* measures how well QC samples are separated from human plasma sample in the first four PC’s. C) R^2^ score between (normalized) abundancies and quantitative measurements. D) Spearman score of (normalized) abundancies with quantitative measurements. E) The WTR score measuring the overall within batch variation with respect to the total variance using the QC samples. F) Δ*R* score measuring the preservation of the rank of features based on their abundancy in the QC samples before and after normalization.

#### Improved separation of QC samples

QC samples (squares in Figure 2) were included in every batch and were expected to separate from the human plasma samples (squares vs circles in Figure 2) in the first four Principle Components (PC) due to overall abundancy differences for several metabolites. Normalization should maintain this separation which was measured by the *QC prediction score* (Figure 3B). *Log-CRMN* conserved QC/plasma separation, with a median *QC prediction score* of 1.00 (1.00) for positive (negative) ion mode, but was less able to reduce batch effects since it had a median *batch prediction score* of 0.76 (0.39) for positive (negative) ion mode respectively. *Log-NOMIS* and *Log-RUV* were better in reducing batch effects, with a median *batch prediction score* of 0.21 (0.21), 0.24 (0.19) for positive (negative) ion mode respectively, but were less able to conserve the separation between QC and human plasma samples, since the median *QC prediction scores* were 0.32 (0.88) and 0.39 (0.94) for positive (negative) ion mode respectively. It is therefore likely that these two methods removed variations other than batch related variation. QC samples were almost perfectly separated from the human plasma sample by *BC-Metchalizer*, *Log-Metchalizer* and *None-Anchor.*

#### Resemblance with quantitative measurements

To further quantify batch effect removal, we calculated the Spearman score and R^2^ score between quantitative plasma concentrations (in μmol/L) and the normalized abundancies of our evaluation set of amino acids and (acyl)carnitines (Methods). To ensure high signal-to-noise ratio’s in the quantitative measurements, we selected only metabolites having a population average concentration above 1 μmol/L. Matching this evaluation set with the annotated features in the untargeted metabolomics data resulted in 16 and 13 metabolites in positive - and negative ion mode, respectively. Figure 3C and D shows both metrics for the investigated normalization methods. Again, for most normalization methods both metrics improved when compared to the raw data (*None-Raw*). *BC-Metchalizer*, *Log-Metchalizer* and *None-Anchor* appeared to perform the best on these metrics with median R^2^ scores of 0.56 (0.55), 0.57 (054), 0.57 (0.47), and median Spearman scores of 0.75 (0.74), 0.74 (0.79), 0.73 (0.71), respectively, for positive (negative) ion mode.

#### Reduced between-batch variation in QC samples

Next, we compared the within-batch variance of the QC samples with respect to the total variance which is expressed by the WTR score (Methods) for each normalization method. WTR scores close to 1 indicate the absence of batch effects. *None-Raw* and *Log-Raw* had low WTR scores and after normalizing these scores increased (Figure 2E). *BC-Metchalizer* and *Log-Metchalizer* scored among the highest on this WTR score. *None-Anchor* had high WTR scores, but since *None-Anchor* uses the QC samples for normalization the WTR scores are biased towards higher values.

#### Preserved feature ranks in QC samples

Removal of variation results in higher WTR scores but potentially removes also variation(s) of interest. Therefore, we investigated whether the ranks of the abundancies of the different features in the QC samples remained the same as in the raw data (expressed as the QC rank differences, Δ*R*, see Methods for details). A lower rank difference indicates that metabolic differences present in the QC samples were conserved after normalization. Figure 2F shows the QC rank differences for each normalization method. These results confirm the previous believe that *Log-NOMIS* and *Log-RUV* also removed non-batch related variations (higher Δ*R*), since they had relatively high Δ*R*’s. *BC-Metchalizer* and *Log-Metchalizer* showed rank differences but were lower than most other competing methods. *None-Anchor* showed high QC rank differences, but this is again the result of the fact that *None-Anchor* uses the QC samples for normalization.

Taken together, *BC-Metchalizer*, *Log-Metchalizer* and *None-Anchor* showed the most consistent improvement across the evaluation metrics.

### Confounder effects of age and sex

To explore confounding effects of age and sex on metabolite abundancies, we developed a regression model with sex as linear covariates and age as a polynomial (*p*=1,2,3) covariate (see Methods). After normalization, we fitted the model parameters for every feature using all control samples present in the eight batches and determined the significance of the coefficients in the regression model (see Methods). Table 1 shows the percentages of (strong) significant coefficients (α = 2.7*e*^−3^) per ion mode and (selected) normalization methods. Our findings suggest that 6-24% of all features showed age dependency when looking at coefficient 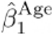 (i.e. the linear term in the model). It is noteworthy that more age-related features were found in the negative ion mode.

**Table 1.**
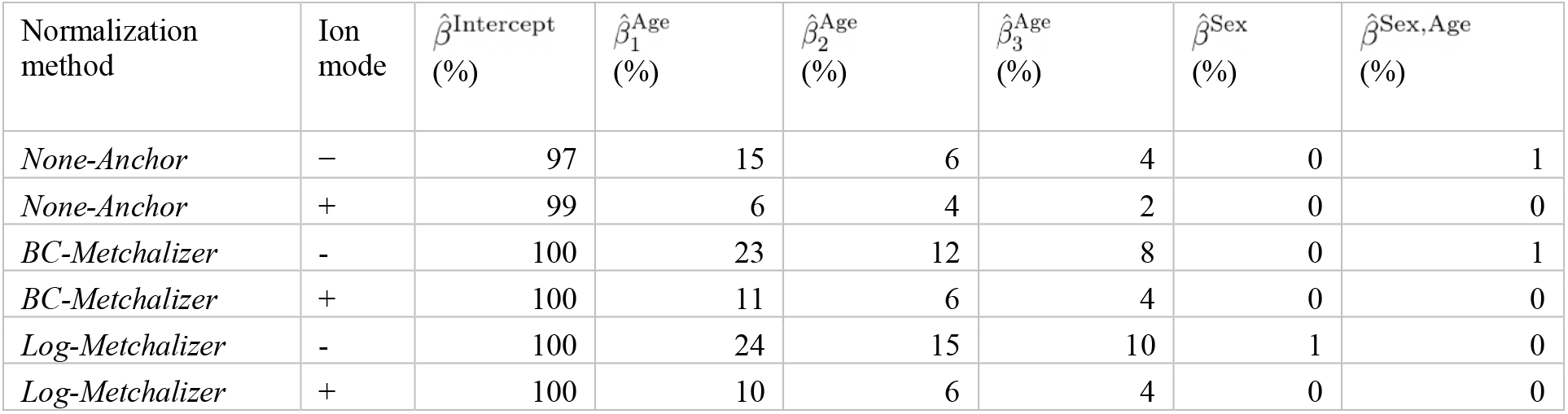
Percentage of strongly significant (α = 2.7*e*^−3^) regression coefficients of the covariates age and sex when using the regression model (Methods) predicting 758 positively - and 583 negatively ionized features, for the different normalization method and ion modes.

Age-dependent metabolites (supplement S3 Table S3), when using normalization by *BC-Metchalizer*, include known IEM biomarkers, such as: guanidinoacetic acid(+), homoarginine(−) and N-acetyltyrosine(−), 2-ketoglutaric acid(−), citrulline(−) and ornithine(−). As an example, we plotted the regression model for guanidinoacetic acid (Figure 3), illustrating that the Z-score for a fixed abundancy depends on age (and slightly on sex at later ages). This also shows a non-linear trend with age. Our analyses showed that more metabolites have significant non-linear trends over age (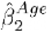 and 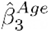 in Table 1). Moreover, age dependent features have the tendency to increase/decrease in abundancy faster for decreasing age, implying that a matching reference population on younger ages seems to be more important (supplement S3 Figure 5).

**Figure 2.**
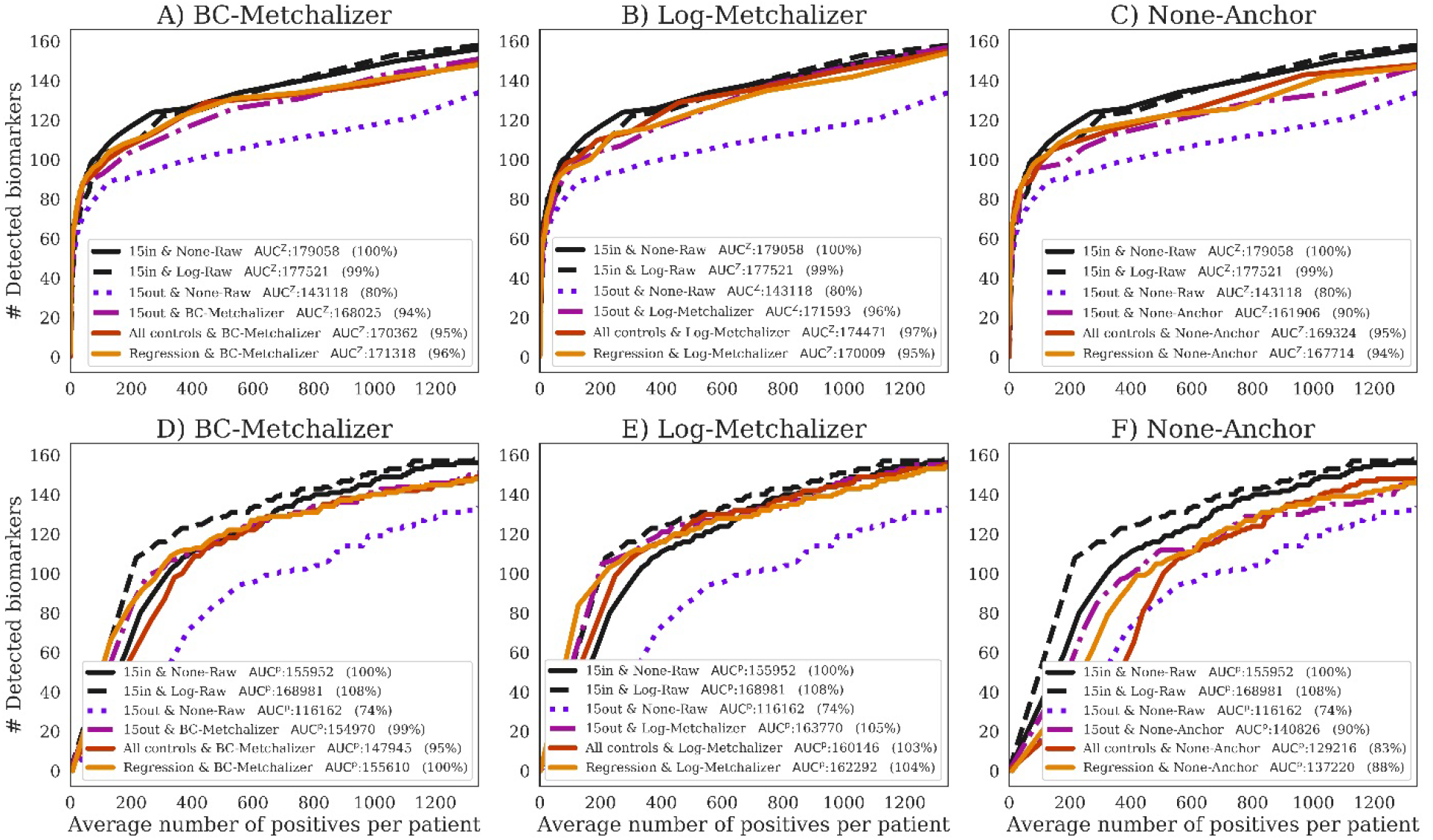
The number of detected expected biomarkers versus the average number of positives per patient. A curve in each (sub)figure was formed by increasing the Z-score or p-values threshold (*Z*_*Zabnormal*_, Methods). The legend indicates (per curve) the methods used to determine Z-scores and how data was normalized, the AUC and AUC expressed as percentage of the 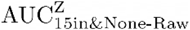. Performances using A) *BC-Metchalizer* using Z-scores, B) *Log-Metchalizer* using Z-scores, C) *None-Anchor* using Z-scores, D) *BC-Metchalizer* using p-values, E) *Log-Metchalizer* using p-values, and F) *None-Anchor* using p-values.

Hardly any significant gender-related features were found (Table 1). When significance on 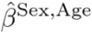 was relaxed (α = 0.05), we found some biomarkers showing an interaction between age and sex, such as: malonic acid(+/−), guanidinoacetic acid(+), homoarginine(−), ornithine(−), sebacic acid(+/−). See supplement S3 Table 4 for a full list.

### Detection of the expected IEM biomarkers

Next we investigated the impact of normalization and using out-of-batch controls on expected biomarker detection in the 45 IEM patients by plotting the number of detected expected biomarkers against the average number of positives features per patient at various Z-score or p-value thresholds (Methods), similar to a Receiver Operator Curve (ROC). Untargeted metabolomics did not allow us to make a distinction between false positives (FP) and true positives (TP), due to unannotated features and even unknown disease related features/biomarkers. Assuming that the majority of the positives per patient are false positives, we used the average number of positives per patient as proxy for the false positives. Improved performance was considered to increase the number of detected expected biomarkers (true positives of which we are certain) while lowering the average number of positives per patient, thereby increasing the Area Under the Receiver Operator Curve (AUC). We decided to take the method that uses 15 within-batch controls and raw abundancies (*15in*&*None-Raw*) as the reference approach, where the performance was expressed as a percentage of this reference AUC, named 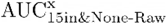. Here indicates if the AUC was created from the average Z-scores or p-values. These p-values were obtained from the Welch’s t-test which tests whether the average Z-score of an expected biomarker or feature across the triplicate significantly differs from the average Z-score of the reference population (Methods).

#### Log-transform improves biomarker detection for p-values

Our first observation is that, when considering the Z-scores, the log-transformed raw abundancies (*15in&Log-Raw*) has an AUC approximately equal to 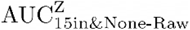 (Figure 4), implying that this transformation hardly affected this performance metric. However, when using the p-values, the log-transformation improved the detection of the expected biomarkers, as 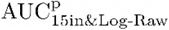 is 8% higher than the 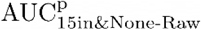 (Figure 4).

**Figure 1.**
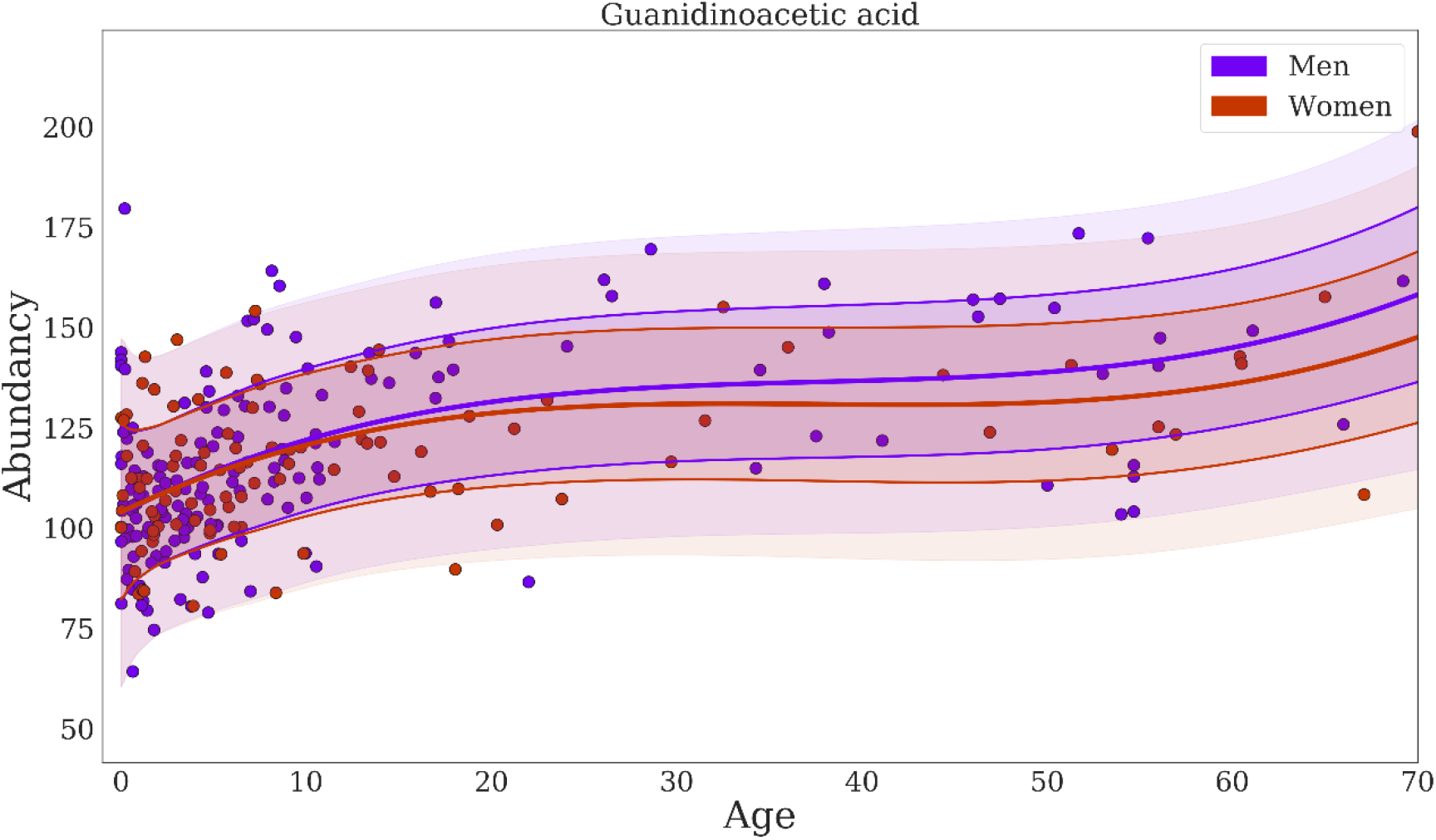
Regression of guanidinoacetic acid when using *BC-Metchalizer* normalized data. The different colors indicate the sex as shown in the legend. The thick red/blue line indicates the average obtained from the fit on all controls for a given sex. The first standard deviation is indicated by the thin(ner) line whereas the second standard deviation ends at the shaded region.

#### Reduced performance with age/sex matched out-of-batch controls

When comparing the performance of using 15 out-of-batch controls (*15out*&*Raw*) to the *15in&Raw* reference, the performance for *15out* was clearly reduced (Figure 4 A), achieving only 80% of the reference 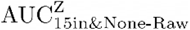. This difference was also present when looking at the p-values, resulting in a clear reduction of the 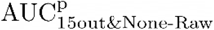 (74%). Hence, potential improved age/sex matching for *15out*, due to the increased number of available controls (supplement S4 Figure 6), did not result in improved performance, most likely due to the dominance of batch effects.

#### Normalization improves performance of age/sex matched out-of-batch controls

After normalizing with *BC-Metchalizer*, *Log-Metchalizer* or *None-Anchor* and using 15 out-of-batch controls (*15out*), the performance increased when compared to *15out*&*None-Raw* (Figure 4 A, B and C), and came close to the 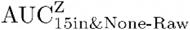; for *BC-Metchalizer* (94%) and *Log-Metchalizer* (96%), while *None-Anchor* stayed behind (90%). Interestingly, when considering biomarker detection performance using the p-values, *BC-Metchalizer* performed on par with *15in&None-Raw* (99%), *Log-Metchalizer* improved over *15in&None-Raw* (105%) and *None-Anchor* stayed behind (90%). *Log-Metchalizer* performed similar to *15in&Log-Raw* (105% and 108%, respectively), indicating that out-of-batch controls can be used instead of in-batch controls to determine reference values.

#### Regression model effectively models age and sex effects

The regression model (*Regression*) slightly improved with respect to *15out* for *BC-Metchalizer* (+2%) and *None-Anchor* (+4%), but not for *Log-Metchalizer* (−1%), see also Figure 4 A, B and C. When considering the p-values,, only *BC-Metchalizer* (+1%) improved but not *None-Anchor* (−2%) and *Log-Metchalizer* (−1%), although these performance differences in all cases were small (Figure 4 D, E and F). Interestingly, when we took all controls to determine the Z-scores (*All controls*, Methods), similar AUC^Z^ performances were observed when compared to *Regression*, i.e. −1% for *BC-Metchalizer* and +2% *Log-Metchalizer* and +1% for *None-Anchor.* When considering the p-values the difference were larger, i.e. −5% for *BC-Metchalizer* and −1% *Log-Metchalizer*, and −5% for *None-Anchor*, suggesting an influence of age- and sex effects on the detection of biomarkers.

## Discussion

Targeted measurements of metabolites in body fluids using various platforms such as HPLC, GC-MS and LC-MS/MS are traditionally applied for laboratory diagnosis of IEM. For each individual metabolite, age- and, sometimes, sex-dependent reference ranges are established using hundreds of reference samples. Untargeted metabolomics is a promising alternative enabling the determination of many metabolites in one analysis. This can speed up the diagnostic process and extends the number of different IEMs that can be screened in a single run. A major drawback of current approaches is that reference samples need to be included in the same experimental batch to ensure proper reference ranges (or Z-score transformations). Some methods do exist that use reference samples measured in different batches (out-of-batch controls) to determine age and sex corrected Z-scores, and they are based on normalizing methods that remove the batch effects. There has not been a comprehensive comparison of the different normalization methods with approaches that use out-of-batch controls, which we have set out in this work. Moreover, we developed a new normalization method, *Metchalizer*, that makes use of isotope-stable internal standards, an approach that has been shown to be useful when mapping specific metabolites to specific internal standards (Körver-Keularts, et al., 2018) which we generalize to all features measured. Because more reference samples are available when using the out-of-batch controls, we additionally propose a regression model that incorporates sex and age effects as (non-linear) covariates. Alltogether, we have shown that our methodology has biomarker detection performances at least similar to using 15 within-batch controls.

Typically, around 20,000 features in both negative and positive mode were detected per batch. When we require a feature to have been measured (and matched) in all eight batches, we retained 598 positive and 773 negative ionized features, respectively. As some normalization methods use a statistical approach (*PQN, Fast Cyclic Loess*), the reduction in features might explain the reduced performance of these methods. In addition, the requirement of features being measured (and matched) across all eight batches also resulted in the loss of some biomarkers, which hampered the performance of all out-of-batch methods with respect to the within-batch methods. As an alternative, we could have made the inclusion of features dependent on fewer batches (for example being present in >5 out of 8 batches). We decided not to do that as this would have resulted in an unequal number of control samples for the different features, leading to inconsistent results between the out-of-batch methods. The availability of more batches could have solved this issue because an equal number of control samples could likely be obtained per feature, even when these features were not present/matched in some batches. It is interesting to note that our proposed normalization method (*Metchalizer*) showed consistent performances when data from various number of batches is being used (supplement Figure S7). Some biomarkers, for example isobutyrylglycine, were only detected in the batches having patients with elevated levels of these specific metabolites. We anticipate that for this kind of biomarkers out-of-batch strategies are not useful since abundancies in (healthy) controls are (very) low, thereby making Z-score calculation unsuitable.

*Anchor* uses an anchor (fixed) samples, measured in all batches, to normalize the features. *Anchor* normalization on none-transformed data performed well when compared to most of the other normalization methods explored, but slightly less than *BC-Metchalizer* and *Log-Metchalizer* when considering the performance metrics Spearman score, R^2^ score, batch prediction score and performance on biomarker detection. We anticipate that the anchor samples may not correlate with all types of variation like, for example, injection volume which is a source of variation at the sample level, whereas the abundancy of the internal standards (used by *Metchalizer*) is directly linked to the injection volume. *Anchor* also assumes that metabolite levels remain constant over time in the anchor samples. As a consequence, if for example storage effects take place, *Anchor* is impeded. The use of *Anchor* may also be less practical because it requires the same anchor samples in every batch. The introduction of a new anchor sample requires an ‘overlapping batch’ containing a set of both the former anchor sample together with the newly introduced anchor samples.

*Metchalizer* exploits the linear relationship between the abundancy of a feature and those of the latent variables that are derived from the partial least squares between the internal standards and the features measured across all samples and capturing the covariance between the standards and the features (Methods). *Metchalizer* assumes that this relationship holds across batches and with that assumption determines (batch) intercepts that correct for the ‘unexplained’ batch/technical variations. Consequently, when such linear relationship between internal standards and features does not exist, the normalization would be fully based on the (batch) intercepts, deteriorating the power of this approach. Alternatively, when batch differences (represented by the intercepts) differ from each other due to biological variations between batches, this will be interpreted as ‘unexplained’ batch/technical variations, and consequently wrongly removed by *Metchalizer*. For this reason, it is important to use randomized control samples in every batch (in terms of age, sex etc) to minimize the possibility of biological variations between batches.

*Log-Metchalizer* log transforms the abundancies before applying *Metchalizer*, whereas *BC-Metchalizer* uses a less strong Box-Cox transformation. The effect of this stronger transformation on most investigated metrics in this study was small, although we did observe that a stronger initial transformation led to improved biomarker detection performances when considering the p-values. *15in*&*None-Raw* had a lower AUC^p^ than *15in*&*Log-Raw* and could therefore also explain the improved performance of *Log-Metchalizer* over *BC-Metchalizer* on this metric. A simulation showed that log-transforming the raw abundancies indeed caused differences in the obtained Z-scores and p-values when compared to the raw abundancies (supplement S10). Positive Z-scores had relatively lower p-values (and vice versa) for log-transformed abundancies and this could therefore explain the improved performance on biomarker detection, since most of the considered biomarkers had positive Z-scores, thus biasing this performance metric. Increasing the number of internal standards did not improve the normalization performance when considering metrics based on the quantitative measurements, although we observed that certain combinations of internal standards improved normalization of specific metabolites (supplement S6). This suggests that *Metchalizer* might be improved by matching features/metabolites with a certain set of internal standards (for example based on retention time).

We were a bit surprised that biomarker detection performance using the Z-scores (AUC^Z^) for the regression model was similar or slightly less than using all controls, as abundancies are known to be dependent on age and sex. One explanation might be that only a subset of the considered (expected) biomarkers have an age and/or sex dependency. Indeed, when we considered only these age-dependent biomarkers (19 biomarker-patient combinations, supplement S3 Table 3), the performance of *Regression* was more improved than *All controls* (supplement S8). However, this set was small, so substantial evidence to support this improvement is lacking. Furthermore, our proposed performance metric assumed that the average number of positives was a proxy for the average number of false positives. Using *Regression* resulted generally in more positives (data not shown), but these were not necessarily merely false positives, which therefore could have affected the performance of *Regression* negatively. Though, when judging biomarker detection using the p-values, we did see that *Regression* slightly outperformed *All controls*.

In conclusion, out of all explored normalization methods, the removal of batch effects was best performed by *Log-Metchalizer.* Fitting our regression model on the corresponding normalized data showed that 10-24% (Table 1) of all considered features were depending on age, underlining the need for using age corrected Z-scores. On average, biomarker detection performance using *Log-Metchalizer* using out-of-batch controls was at least similar to the best performing *Log-Raw* approach when using the 15 within-batch controls (*15in&Log-Raw*). We anticipate that the success of *Metchalizer* and age- and sex correcting strategies such as our regression model depend on three factors: 1) a feature of interest being measured in a number of other batches (not necessarily all), 2) batch effects containing (only) technical variations, and 3) abundancies being affected by age or other covariates (the presence of an effect-size). Together our proposed approach opens new opportunities to improve abnormality detection, especially for age-dependent features/biomarkers.

## Author contribution

Ramon Bonte performed all the experimental work and developed the chromatographic- and mass spectrometric method. Compound identification was also done by him. Michiel Bongaerts designed the statistical models, the computational framework and analyzed the data. The manuscript was written by Michiel Bongaerts, Henk Blom and George Ruijter. Serwet Demirdas and Ed Jacobs contributed in establishing the IEM database used in this study, and actively contributed in giving feedback on the methods. Marcel Reinders contributed to in-depth reviewing of the manuscript, all analytical methods and suggested adjustments to initial work. Esmee Oussoren, Ans van der Ploeg, Margreet Wagenmakers and Robert Hofstra provided resources. The research was under supervision of George Ruijter.

## Data and Code availability

The regression model, *Best Correlated Internal Standard, PQN, Anchor* and *Metchalizer(Log)* were developed in Python and are available at https://github.com/mbongaerts/Metchalizer. The code developed for merging the batches can also be found here. The Progenesis QI processed data for all 8 batches is available at https://github.com/mbongaerts/Metchalizer/Data. We removed the patient samples for privacy reasons.

