## Supplementary data for "Screening for inborn errors of metabolism using untargeted metabolomics and out-of-batch controls" (from another version of this preprint: version 1, 2020-04-14)

### S1. Inter- and intra-batch Coefficients of Variation

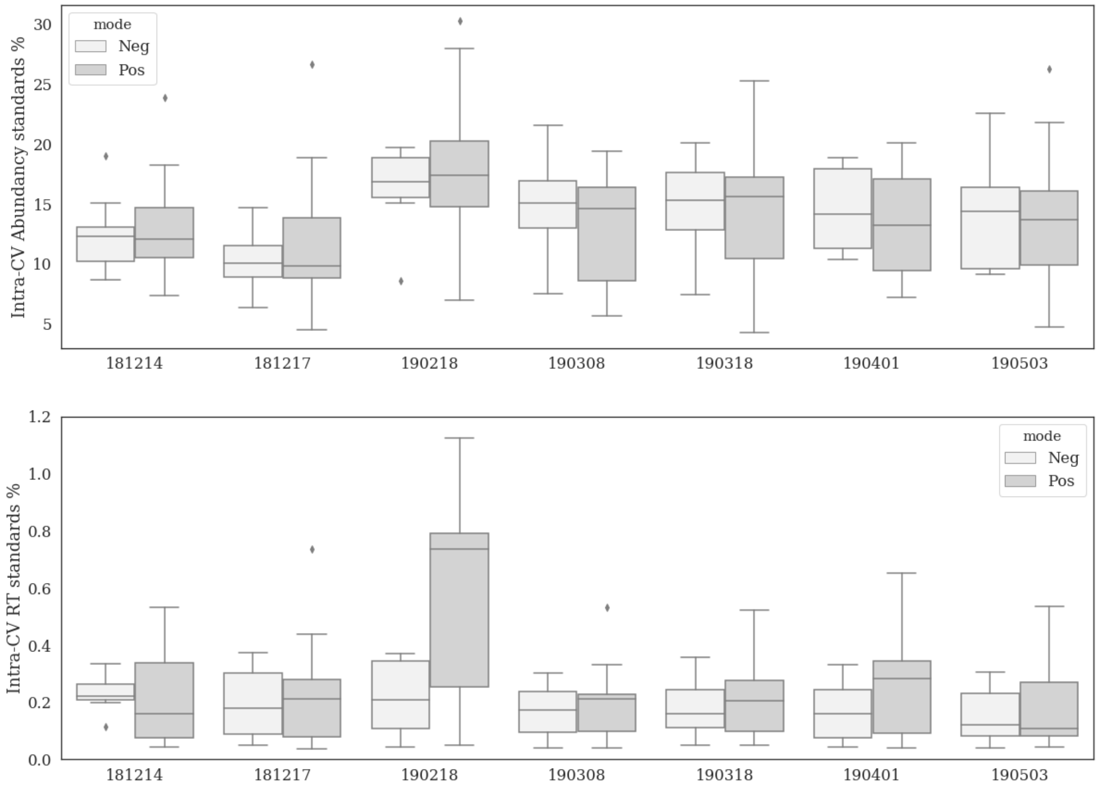

Figure 1. Boxplots per batch and ion mode for intra-batch Coefficients of Variation (CV) for all standards. Top) Intra-batch CVs for abundancies determined by using Xcalibur 4.0. Bottom) Intra-batch CVs for retention time.

Table 1. Inter-batch Coefficients of Variation per standard based on 8 batches. Note, that some components were detected in both ion modi but the best CV was considered for quality control. Peak areas were determined using Xcalibur 4.0.

| **Standard** | **Ion mode** | **Inter batch CV (%)** |
|---|---|---|
| 3,3-dimethylglutaric acid | - | 24 |
| 3,3-dimethylglutaric acid | + | 71 |
| 5-bromotryptophane | - | 18 |
| 5-bromotryptophane | + | 22 |
| Acetylcarnitine D2 | + | 29 |
| Carnitine D3 | + | 22 |
| Glycochenodeoxycholic Acid D4 | - | 22 |
| Hexadecanoylcarnitine D3 | + | 35 |
| Hexanoylcarnitine D3 | + | 27 |
| Isoleucine D10 | - | 38 |
| Isoleucine D10 | + | 27 |
| Methylmalonic Acid D3 | - | 49 |
| Ornithine D6 | + | 26 |
| Phenylalanine D5 | - | 38 |
| Phenylalanine D5 | + | 28 |
| Tetradecanoylcarnitine D3 | + | 28 |
| Thymidine 13C | - | 18 |
| Thymidine 13C | + | 50 |
| Tyrosine D4 | - | 30 |
| Tyrosine D4 | + | 27 |
| Uracil-1,3N15 | - | 19 |
| Uracil-1,3N15 | + | 20 |
| Uridine D2 | - | 27 |
| Uridine D2 | + | 54 |
| Valine D8 | + | 40 |

### S2. Performances normalization methods

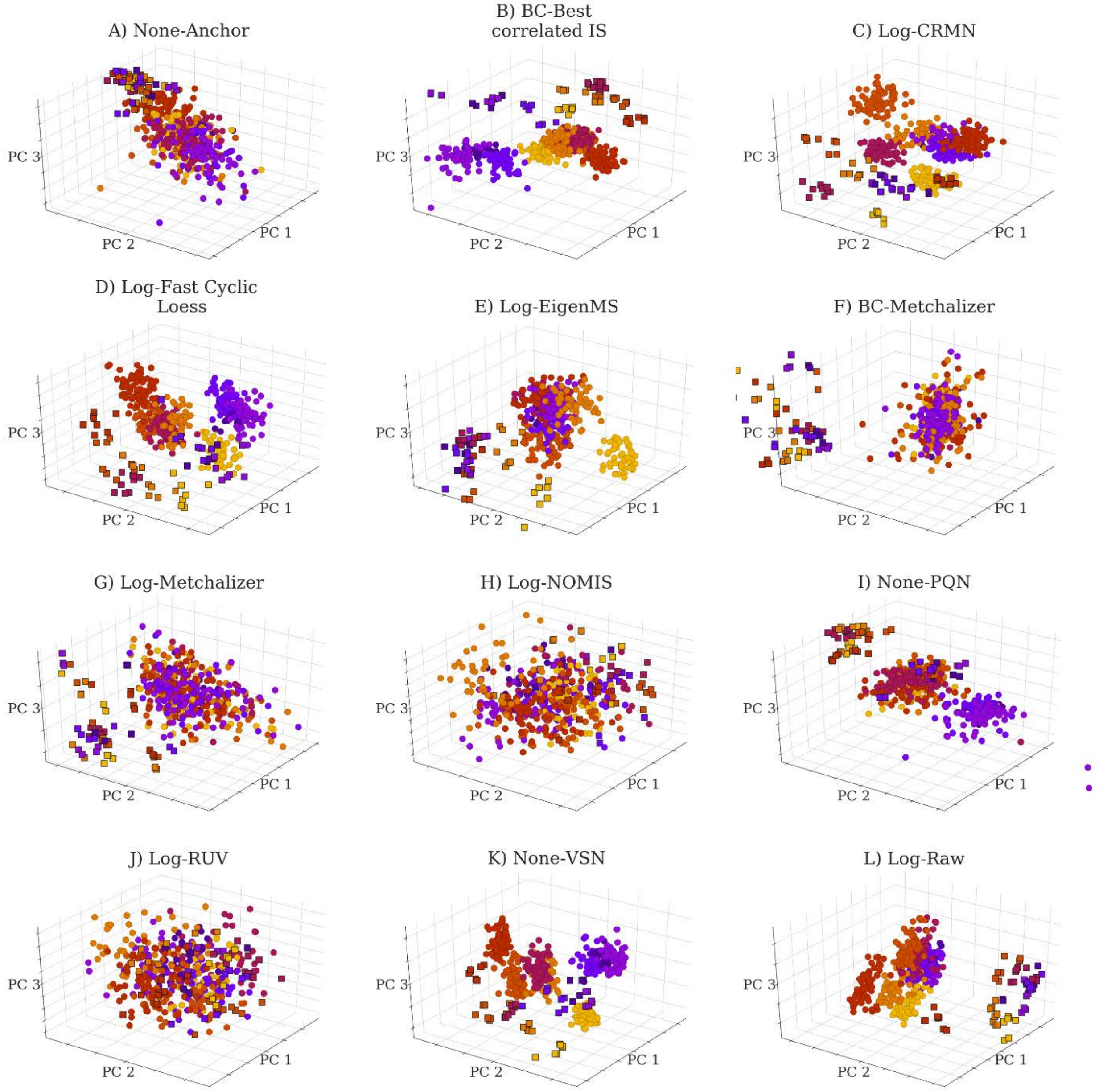

Figure 2. PCA plots for a given normalization method. Each batch is indicated with an unique color. PCA was performed only the 758 features (excluding standards) in **positive** ion mode. The squares indicate QC samples whereas the dots indicate human plasma's.

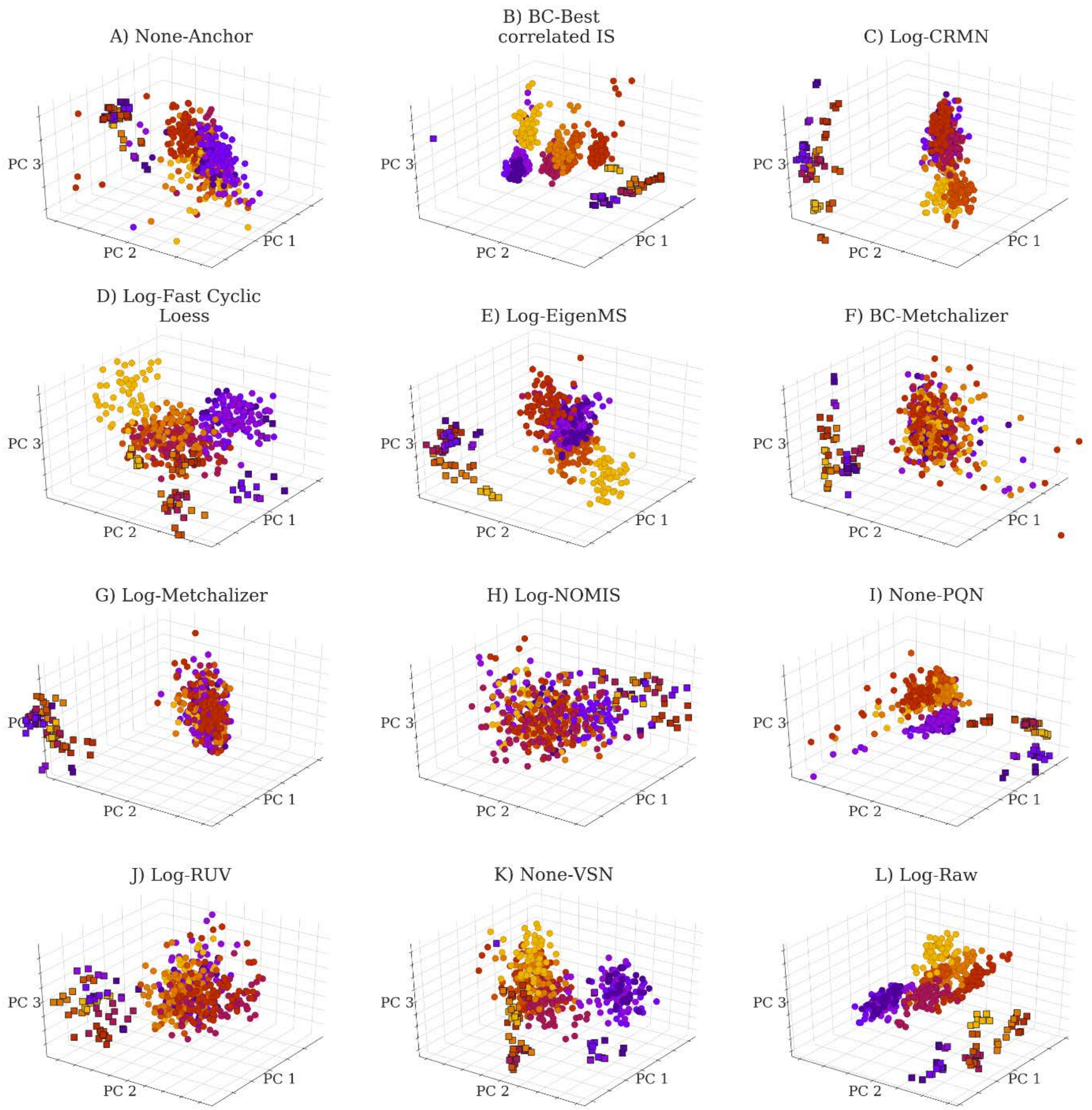

Figure 3. PCA plots for a given normalization method. Each batch is indicated with an unique color. PCA was performed only the 583 features (excluding standards) in **negative** ion mode. The squares indicate QC samples whereas the dots indicate human plasma's.

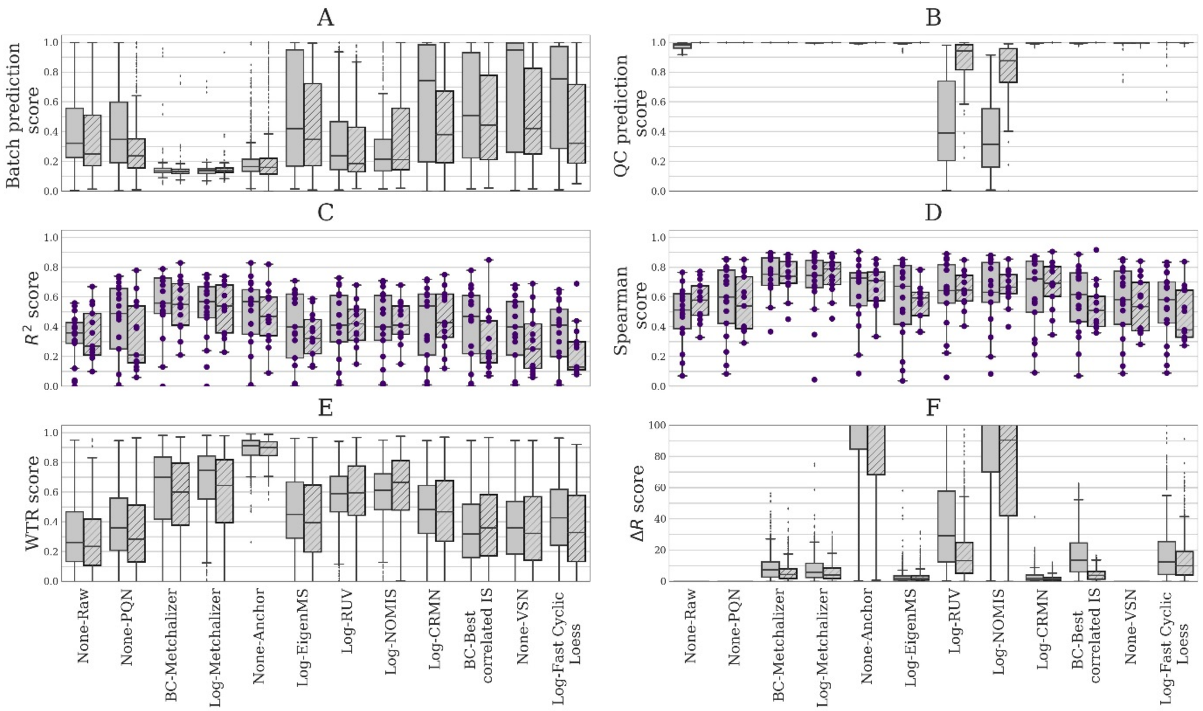

Figure 4. Six different performance metrics for batch effect removal (see Methods for more details). Data from positive – or negative ion mode is indicated by plain, stripped boxplots respectively. A) *Batch prediction score* measures the presence of batch effects in the first four PC's. B) *QC prediction score* measures how well QC samples are separated from human plasma sample in the first four PC's. C) $R^2$ between (normalized) abundancies and quantitative measurements. D) Spearman correlation of (normalized) abundancies with quantitative measurements. E) The WTR scores measuring the overall within batch variation with respect to the total variance. F) Rank difference ($\Delta R$) measuring the rank based on abundancy of features in the QC samples before and after normalization.

Figure 4. shows six metrics to evaluate batch effect removal (Methods). Table 2 shows for four of these metrics the median score for each normalization method and ion mode. We only included normalization methods in the main text for which in both ion modi the average score was >= 0.5. Note that the WTR-scores for *None-Anchor* are biased towards higher values since *None-Anchor* used the QC-samples for normalization.

Table 2. Median scores for four metrics per normalization method and ion mode. The last column is the average of the former columns. Note, that we modified the batch prediction score such that higher values correspond with improved normalization performances.

| **Method** | **Mode** | **$R^2$** | **Spearman** | **WTR score** | **1- Median batch prediction score** | **Mean score** |
|---|---|---|---|---|---|---|
| *None-Anchor* | + | 0.57 | 0.73 | 0.91 | 0.83 | 0.76 |
| *None-Anchor* | - | 0.47 | 0.71 | 0.9 | 0.84 | 0.73 |
| *Log-Metchalizer* | + | 0.57 | 0.74 | 0.75 | 0.86 | 0.73 |
| *BC-Metchalizer* | + | 0.56 | 0.75 | 0.7 | 0.87 | 0.72 |
| *Log-Metchalizer* | - | 0.54 | 0.79 | 0.65 | 0.86 | 0.71 |
| *BC-Metchalizer* | - | 0.55 | 0.74 | 0.6 | 0.87 | 0.69 |
| *Log-NOMIS* | - | 0.41 | 0.67 | 0.67 | 0.79 | 0.64 |
| *Log-RUV* | - | 0.42 | 0.65 | 0.6 | 0.81 | 0.62 |
| *Log-NOMIS* | + | 0.4 | 0.63 | 0.61 | 0.79 | 0.61 |
| *Log-RUV* | + | 0.41 | 0.64 | 0.59 | 0.76 | 0.6 |
| *Log-CRMN* | - | 0.43 | 0.69 | 0.47 | 0.62 | 0.55 |
| *Log-EigenMS* | + | 0.4 | 0.67 | 0.45 | 0.58 | 0.52 |
| *None-PQN* | + | 0.49 | 0.6 | 0.36 | 0.65 | 0.52 |
| *Log-CRMN* | + | 0.54 | 0.72 | 0.48 | 0.26 | 0.5 |
| *Log-EigenMS* | - | 0.32 | 0.59 | 0.39 | 0.65 | 0.49 |
| *BC- Best Correlated IS* | + | 0.47 | 0.62 | 0.32 | 0.49 | 0.48 |
| *None-Raw* | - | 0.27 | 0.58 | 0.23 | 0.75 | 0.46 |
| *None-Raw* | + | 0.36 | 0.51 | 0.26 | 0.68 | 0.45 |
| *None-PQN* | - | 0.21 | 0.54 | 0.28 | 0.76 | 0.45 |
| *Log-Raw* | - | 0.32 | 0.59 | 0.25 | 0.57 | 0.43 |
| *None-VSN* | - | 0.25 | 0.54 | 0.32 | 0.58 | 0.42 |
| *Log- Fast Cyclic Loess* | + | 0.41 | 0.58 | 0.43 | 0.24 | 0.42 |
| *BC- Best Correlated IS* | - | 0.22 | 0.51 | 0.36 | 0.56 | 0.41 |
| *Log-Fast Cyclic Loess* | - | 0.13 | 0.38 | 0.33 | 0.68 | 0.38 |
| *Log-Raw* | + | 0.32 | 0.51 | 0.26 | 0.31 | 0.35 |
| *None-VSN* | + | 0.4 | 0.58 | 0.36 | 0.05 | 0.35 |

### S3. Regression model

Table 3. Significant age related metabolites (p-value $\hat{\beta}_1^{\text{Age}}$ < 2.7e-3 ) for *BC-Metchalizer* normalized data. Slashes in the names indicate that chromatographic separation of isomers was not possible. Sign $\hat{\beta}_1^{\text{Age}}$ indicates if this coefficient is positive (up) or negative (down).

| **Metabolite** | **Ion mode** | **Sign** $\hat{\beta}_1^{\text{Age}}$ | **P-value** $\hat{\beta}^{\text{Intercept}}$ | **P-value** $\hat{\beta}_1^{\text{Age}}$ | **P-value** $\hat{\beta}_2^{\text{Age}}$ | **P-value** $\hat{\beta}_3^{\text{Age}}$ | **P-value** $\hat{\beta}^{\text{Sex}}$ | **P-value** $\hat{\beta}^{\text{Sex,Age}}$ |
|---|---|---|---|---|---|---|---|---|
| 1-Methyladenosine (1) | + | down | 0 | 2.9e-06 | 5.5e-05 | 1.1e-03 | 1.1e-01 | 3.3e-01 |
| 2-Aminoadipic acid | + | down | 0 | 1.8e-06 | 4.0e-05 | 7.4e-04 | 7.5e-01 | 1.5e-01 |
| 2-Aminoadipic acid | - | down | 0 | 4.6e-07 | 4.3e-06 | 1.1e-04 | 8.5e-01 | 1.5e-01 |
| 2-Hydroxybutyric acid | - | down | 0 | 1.7e-03 | 1.3e-02 | 3.1e-02 | 4.7e-01 | 3.9e-01 |
| 2-Ketoglutaric acid | - | down | 0 | 7.1e-06 | 1.5e-05 | 8.5e-05 | 7.6e-01 | 4.9e-01 |
| 3-Methoxytyrosine | + | down | 0 | 2.6e-10 | 7.6e-05 | 9.2e-03 | 7.8e-01 | 6.3e-01 |
| 4-Pyridoxic acid | - | down | 0 | 2.5e-05 | 2.2e-03 | 2.5e-02 | 1.0e-01 | 6.9e-02 |
| 5-Methylcytosine | + | down | 0 | 8.9e-06 | 5.9e-04 | 4.8e-03 | 2.6e-01 | 4.5e-01 |
| 6-Deoxygalactose | - | up | 0 | 1.7e-10 | 4.5e-09 | 1.2e-07 | 8.0e-01 | 2.8e-02 |
| Acetoacetic acid / Succinic acid semialdehyde | + | down | 0 | 4.3e-04 | 1.2e-02 | 6.1e-02 | 7.3e-01 | 2.8e-01 |
| Adenine | + | down | 0 | 2.1e-03 | 1.6e-02 | 6.6e-02 | 4.4e-01 | 4.0e-01 |
| Alanine / β-Alanine / Sarcosine | - | down | 0 | 1.6e-03 | 6.2e-01 | 6.5e-01 | 7.5e-01 | 3.2e-01 |
| C2 \| Acetylcarnitine | + | down | 0 | 9.2e-04 | 6.8e-03 | 2.5e-02 | 3.5e-02 | 2.0e-01 |
| C3 \| Propionylcarnitine | + | down | 0 | 2.2e-03 | 3.8e-03 | 1.4e-02 | 8.1e-01 | 2.0e-01 |
| C8:1 \| Octenoylcarnitine | + | down | 0 | 1.4e-03 | 3.6e-02 | 2.2e-01 | 5.1e-01 | 4.9e-01 |
| C8DC \| Suberylcarnitine | + | down | 0 | 3.2e-07 | 1.9e-05 | 2.3e-04 | 4.4e-01 | 6.3e-02 |
| Chenodeoxycholic acid | - | up | 0 | 7.2e-07 | 1.1e-01 | 6.1e-01 | 6.0e-02 | 4.0e-01 |
| Citrulline | - | up | 0 | 2.3e-04 | 1.6e-02 | 4.7e-02 | 2.6e-01 | 5.3e-02 |
| Creatine | + | down | 0 | 1.1e-03 | 7.3e-01 | 4.4e-01 | 8.9e-01 | 6.1e-01 |
| Creatinine | + | up | 0 | 2.9e-13 | 1.7e-03 | 1.4e-01 | 8.8e-01 | 2.0e-01 |
| Dehydro-epiandrosteronsulfaat (dheas) | - | up | 0 | 2.9e-11 | 3.9e-02 | 8.8e-01 | 9.0e-01 | 9.5e-02 |
| Dimethylarginine (sdma + adma) | - | down | 0 | 5.3e-07 | 2.3e-04 | 4.3e-03 | 9.5e-02 | 4.8e-01 |
| Dimethylarginine (sdma + adma) | + | down | 0 | 4.6e-09 | 5.5e-05 | 3.1e-03 | 1.4e-01 | 2.4e-01 |
| Formyl-l-methionyl peptide | - | down | 0 | 2.7e-05 | 4.1e-04 | 4.8e-03 | 7.3e-01 | 4.8e-01 |
| Glucuronic acid / Galacturonic acid / Gluconic acid | - | down | 0 | 2.3e-03 | 2.5e-02 | 1.9e-01 | 6.2e-01 | 3.3e-01 |
| Glutamic acid / N-Methyl-D-Aspartic acid | + | down | 0 | 6.0e-04 | 2.6e-03 | 1.1e-02 | 6.0e-01 | 5.7e-01 |
| Glycocholic acid | - | down | 0 | 2.3e-03 | 2.5e-01 | 8.0e-01 | 4.3e-01 | 5.0e-01 |
| Glycocholic acid | + | down | 0 | 1.4e-03 | 1.5e-01 | 5.5e-01 | 4.8e-01 | 5.5e-01 |
| Guanidinoacetic acid | + | up | 0 | 4.6e-05 | 1.1e-01 | 3.7e-01 | 5.9e-01 | 2.1e-02 |
| Hexose-1 / 6-phosphate | + | down | 0 | 1.2e-03 | 6.7e-01 | 5.0e-01 | 6.0e-01 | 1.7e-01 |
| Homoarginine | - | up | 0 | 2.5e-08 | 3.6e-03 | 1.4e-01 | 1.9e-01 | 4.3e-02 |
| Hydantoin-5-propionic acid | + | down | 0 | 2.2e-05 | 9.8e-04 | 1.2e-02 | 5.9e-01 | 2.0e-01 |
| Indoxylsulfuric acid | - | up | 0 | 2.3e-03 | 1.9e-02 | 4.8e-02 | 1.3e-01 | 7.5e-01 |
| Kynurenin | + | down | 0 | 1.7e-08 | 5.2e-05 | 2.1e-03 | 5.8e-01 | 9.3e-01 |
| N-Acetylaspartic acid | - | down | 0 | 1.6e-11 | 9.1e-06 | 1.9e-03 | 3.5e-01 | 6.4e-01 |
| N-Acetyltyrosine | - | down | 0 | 4.2e-05 | 6.0e-04 | 8.6e-03 | 3.0e-01 | 8.4e-01 |
| Ornithine | - | up | 0 | 3.8e-04 | 2.2e-02 | 5.5e-02 | 1.9e-01 | 3.2e-02 |
| Pantothenic acid | + | down | 0 | 0 | 3.4e-10 | 6.4e-07 | 1.4e-01 | 3.1e-01 |

| | | | | | | | | |
|---|---|---|---|---|---|---|---|---|
| Pantothenic acid | - | down | 0 | 1.0e-15 | 2.9e-09 | 5.4e-06 | 1.4e-01 | 3.5e-01 |
| Pseudouridine | + | down | 0 | 5.5e-06 | 8.3e-05 | 9.5e-04 | 2.0e-01 | 1.8e-01 |
| Raffinose / Hex3 | - | down | 0 | 2.4e-05 | 2.1e-03 | 2.6e-02 | 2.4e-01 | 7.6e-01 |
| Raffinose / Hex3 | + | down | 0 | 3.1e-06 | 1.6e-04 | 2.7e-03 | 8.2e-02 | 6.0e-01 |
| Ribose / Xylose / Arabinose | - | down | 0 | 2.9e-07 | 1.8e-04 | 5.1e-03 | 7.4e-01 | 2.5e-01 |
| Sialic acid | + | down | 0 | 3.9e-06 | 1.1e-03 | 2.4e-02 | 2.7e-01 | 4.4e-01 |
| Sialic acid | - | down | 0 | 2.5e-10 | 1.2e-05 | 2.0e-03 | 3.9e-01 | 8.2e-01 |
| Sucrose / Lactose / Maltose | + | down | 0 | 4.7e-07 | 2.9e-04 | 6.3e-03 | 6.2e-01 | 4.6e-01 |
| Theophylline /Paraxanthine | + | up | 0 | 2.5e-03 | 7.2e-01 | 8.7e-01 | 2.9e-01 | 7.1e-02 |

Table 4. Significant (p-value $\hat{\beta}^{\text{Sex,Age}}$ < 0.05) interaction of age and sex related metabolites for *BC-Metchalizer* normalized data. An “up” in column Sign $\hat{\beta}^{\text{Sex,Age}}$ indicates that the metabolite was increased in women and vice versa. Slashes in the names indicate that chromatographic separation was not possible.

| **Metabolite** | **Ion mode** | **Sign** $\hat{\beta}^{\text{Sex,Age}}$ | **P-value** $\hat{\beta}^{\text{Intercept}}$ | **P-value** $\hat{\beta}_1^{\text{Age}}$ | **P-value** $\hat{\beta}_2^{\text{Age}}$ | **P-value** $\hat{\beta}_3^{\text{Age}}$ | **P-value** $\hat{\beta}^{\text{Sex}}$ | **P-value** $\hat{\beta}^{\text{Sex,Age}}$ |
|---|---|---|---|---|---|---|---|---|
| 4-Hydroxyphenylpyruvic acid | - | up | 0 | 5.6e-01 | 4.4e-01 | 6.2e-01 | 7.6e-01 | 4.9e-02 |
| 5-Oxoproline | + | down | 0 | 7.6e-01 | 8.4e-01 | 8.3e-01 | 1.3e-01 | 3.4e-02 |
| 6-Deoxygalactose | - | down | 0 | 1.7e-10 | 4.5e-09 | 1.2e-07 | 8.0e-01 | 2.8e-02 |
| Betaine | + | down | 0 | 5.6e-01 | 8.8e-01 | 7.8e-01 | 8.8e-01 | 3.9e-03 |
| C6 \| Hexanoylcarnitine | + | up | 0 | 7.0e-03 | 2.6e-02 | 5.1e-02 | 1.1e-02 | 1.0e-03 |
| Glutamine + Glutamic acid / N-Methyl-D-Aspartic acid | - | down | 0 | 7.0e-02 | 6.3e-02 | 1.0e-01 | 1.9e-01 | 2.2e-02 |
| Glutaric acid | + | up | 0 | 7.4e-01 | 6.0e-01 | 6.9e-01 | 3.5e-01 | 1.3e-02 |
| Guanidinoacetic acid | + | down | 0 | 4.6e-05 | 1.1e-01 | 3.7e-01 | 5.9e-01 | 2.1e-02 |
| Homoarginine | - | down | 0 | 2.5e-08 | 3.6e-03 | 1.4e-01 | 1.9e-01 | 4.3e-02 |
| Maleic acid | - | up | 0 | 9.0e-01 | 2.7e-01 | 2.0e-01 | 3.0e-01 | 7.6e-03 |
| Malonic acid | + | up | 0 | 4.7e-01 | 3.2e-01 | 4.3e-01 | 1.4e-01 | 8.3e-03 |
| Malonic acid | - | up | 0 | 6.0e-01 | 1.7e-01 | 1.6e-01 | 4.3e-01 | 2.5e-02 |
| N-Phenylacetylglutamic acid | - | up | 0 | 6.9e-01 | 7.4e-01 | 7.1e-01 | 7.0e-01 | 1.7e-02 |
| Nicotinic acid / Picolinic acid | + | up | 0 | 4.2e-02 | 1.9e-01 | 2.2e-01 | 8.6e-01 | 1.5e-02 |
| Ornithine | - | down | 0 | 3.8e-04 | 2.2e-02 | 5.5e-02 | 1.9e-01 | 3.2e-02 |
| Sebacic acid | - | up | 0 | 7.6e-01 | 4.2e-01 | 3.9e-01 | 3.3e-01 | 4.9e-02 |
| Sebacic acid | + | up | 0 | 8.6e-01 | 6.3e-01 | 6.2e-01 | 9.5e-02 | 9.7e-03 |
| Succinic acid | - | up | 0 | 2.9e-01 | 1.3e-01 | 1.6e-01 | 3.3e-01 | 4.1e-02 |
| Trigonelline | + | down | 0 | 2.4e-01 | 7.7e-01 | 8.8e-01 | 2.1e-02 | 4.7e-03 |
| Tryptophan | - | down | 0 | 1.6e-01 | 9.4e-02 | 1.1e-01 | 1.9e-01 | 4.4e-02 |

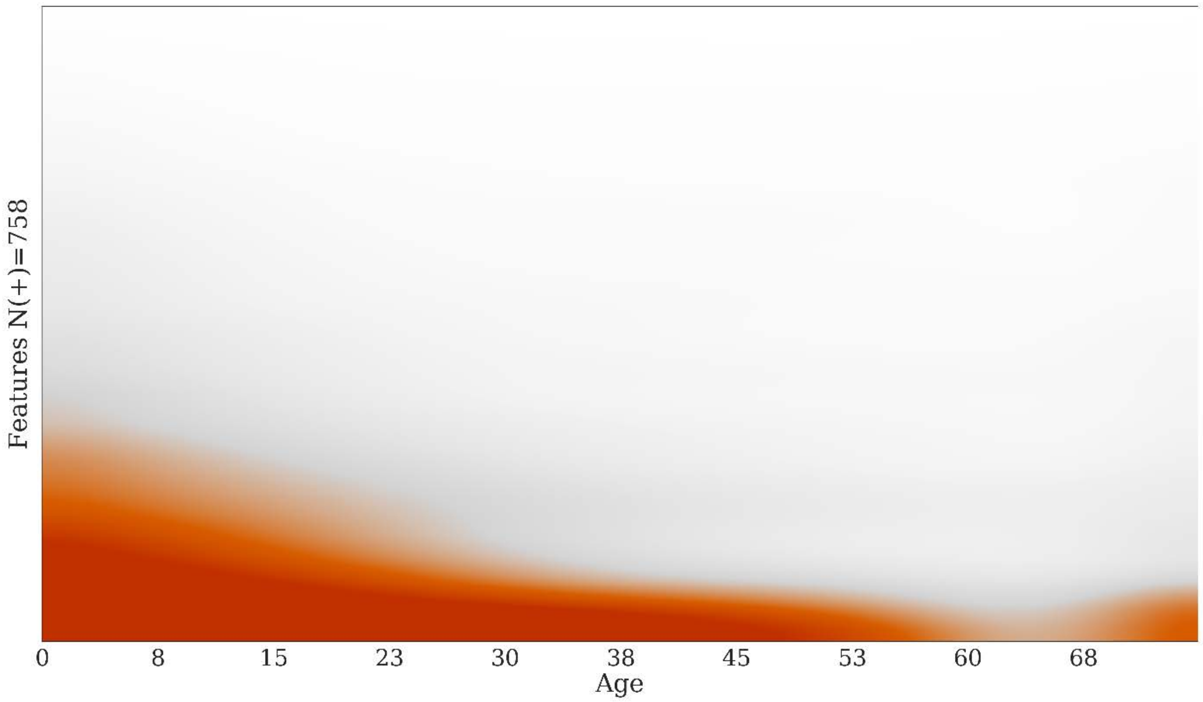

Figure 5. By taking the derivative of the regression model with respect to $x_i^{\mathrm{Age}}$ we obtained the slope at a given age. This figure was obtained by plotting these slopes for all features (in positive ion mode and excluding internal - and external standards) where slopes were first divided by the median abundancy to obtain comparable numbers where after the absolute value was taken. Red and orange area's indicates larger values of the slopes. Colors were also smoothed. The data was normalized using *BC-Metchalizer*.

### S4. Similarity of controls with patients for different methods

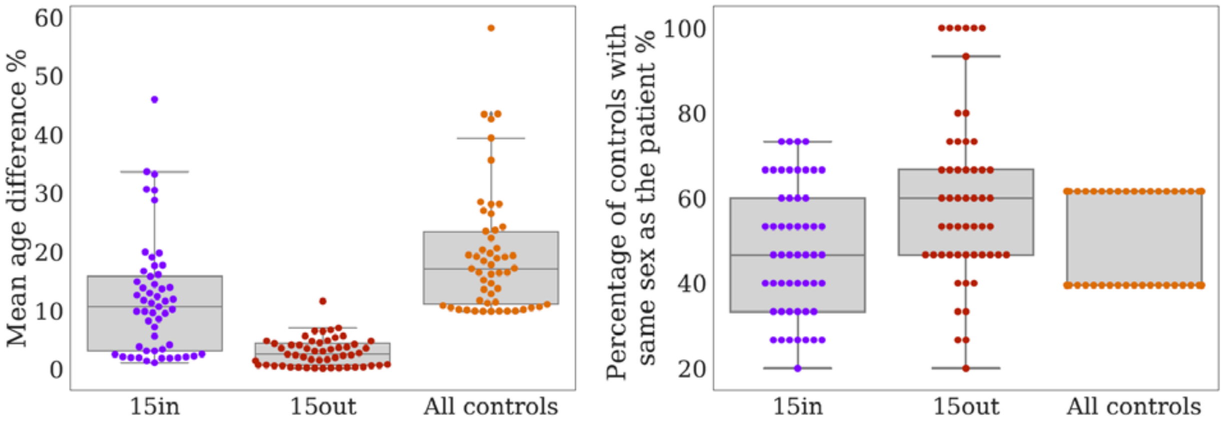

Figure 6. Left) Similarity in age between patient and controls for every method. Mean age difference was determined by calculating the difference in age of the patient with the controls divided by the age of the patient (times 100 %) where after the mean was taken. Every dot represent this mean for one patient. Right ) Similarity in sex between patient and controls for every method. Every dot represents a patient.

### S5. Z-score table per IEM and method

Table 5. The Z-scores obtained from *Raw&15in* , *Metchalizer&Regression* and *Anchor&Regression* per biomarker and IEM. The stars on the Z-scores indicate that Welch's t-test p-values < 0.0. The amount of unique patients is indicated by *N* next to the name of the IEM.

| **Biomarker (expected Z-score sign) (ion mode)** | **None-Raw & 15in** | **BC-Metchalizer & Regression** | **None-Anchor & Regression** |
|---|---|---|---|
| **Aminoacylase I deficiency N=1** | | | |
| N-Acetylglycine (up) (-) | 9.3* | 4.1* | 7.4* |
| N-Acetylglycine (up) (+) | 5.3* | 1.3* | 3.7* |
| **Argininemia N=1** | | | |
| 4-Guanidinobutyric acid (up) (+) | 25.6* | 11.8* | 24.6* |
| Arginine (up) (-) | 6.0* | 3.9* | 6.8* |
| Arginine (up) (+) | 3.7* | 2.8* | 5.4* |
| Glutamine + Glutamic acid / N-Methyl-D-Aspartic acid (up) (-) | 0.4 | 0.3 | 1 |
| Guanidinoacetic acid (down) (+) | 3.8* | 2.4* | 2.4* |
| Homoarginine (up) (-) | 13.9* | 6.9* | 15.0* |
| Uridine (up) (-) | 4.0* | 3.7* | 3.7* |
| **Argininosuccinic aciduria N=3** | | | |
| Arginine (down) (-) | -0.9*, -0.2, 2.0* | -1.0*, 0.2, 1.1* | -1.0*, 0.9, 0.7* |
| Arginine (down) (+) | -1.0*, -0.1, 2.0* | -1.3*, 0.2, 1.3* | -1.3*, 1.2*, 1.2* |
| Citrulline (up) (-) | 30.7*, 19.6*, 17.9* | 9.1*, 9.4*, 8.0* | 20.1*, 17.3*, 11.6* |
| Citrulline (up) (+) | 19.9*, 18.5*, 12.9* | 7.0*, 10.6*, 8.2* | 16.2*, 16.7*, 9.9* |
| Glutamine + Glutamic acid / N-Methyl-D-Aspartic acid (up) (-) | 2.7*, 0.4, 1.3* | 1.5*, 1.2, 1.5* | 1.1*, 1.7*, 1.8* |
| Homocitrulline (up) (-) | 4.8*, 2.2*, 4.3* | 4.9*, 1.7*, 2.0* | 10.4*, 1.7*, 2.1* |
| Homocitrulline (up) (+) | 4.3*, 2.1*, 3.8* | 4.4*, 1.2*, 1.5* | 11.0*, 1.5*, 2.3* |
| Uridine (up) (-) | 0.5, -0.2, 5.3* | 0.2, -0.7*, 4.3* | 0.4*, -0.6*, 4.4* |
| **Beta-ketothiolase deficiency N=2** | | | |
| 2-Methylacetoacetic acid (up) (+) | 0.4, -1.1* | 0.9, 0.7 | 0.4, 0.6 |
| **Carbamoyl Phosphate Synthetase deficiency N=2** | | | |
| 2-Ketoglutaric acid (up) (-) | -1.1*, -0.8 | -1.5*, -1.1 | -0.6*, -0.5 |
| Glutamine + Glutamic acid / N-Methyl-D-Aspartic acid (up) (-) | 3.6*, -0.5 | 2.4*, 0.1 | 3.9*, 0.4 |
| **Citrullinemia type I N=1** | | | |
| Arginine (down) (-) | 0.9* | 0.8 | -0.2 |

| | | | |
|---|---|---|---|
| Arginine (down) (+) | 0.9 | 1 | 0.4 |
| Citrulline (up) (-) | 153.0* | 52.1* | 187.0* |
| Citrulline (up) (+) | 108.7* | 47.6* | 109.0* |
| Glutamine + Glutamic acid / N-Methyl-D-Aspartic acid (up) (-) | -0.5 | -1.1 | -0.8* |
| Uridine (up) (-) | 1.5* | 1.2* | 0.5 |
| **Glutamate formiminotransferase deficiency N=1** | | | |
| Formiminoglutamic acid (up) (+) | 2662.6* | 136.7* | 849.1* |
| **Glutaric aciduria I N=2** | | | |
| C5DC \| Glutarylcarnitine (up) (+) | 317.5*, 324.0* | 28.0*, 36.6* | 293.6*, 454.9* |
| Glutaric acid (up) (+) | 0.9, 2.2* | 0.5, 2.3* | 0.8, 1.8* |
| **Glutaric aciduria II N=2** | | | |
| Adipic acid (1) (up) (-) | 1.5*, 8.0* | 1.3*, 5.2* | 1.3*, 6.8* |
| C10 \| Decanoylcarnitine (up) (+) | 62.5*, 58.6* | 23.0*, 20.3* | 63.4*, 39.1* |
| C12 \| Dodecanoylcarnitine (up) (+) | 0.9, 69.1* | 0.5*, 23.4* | 0.8, 34.9* |
| C14 \| Tetradecanoylcarnitine (up) (+) | 149.4, 112.0* | 23.1, 16.1* | 94.5, 71.6* |
| C14:1 \| Tetradecenoylcarnitine (up) (+) | 97.6, 70.2* | 19.3*, 18.0* | 104.6, 92.3* |
| C16 \| Hexadecanoylcarnitine (up) (+) | 6.7, 10.6* | 4.8, 3.2* | 6.3, 4.2* |
| C16:1 \| Hexadecenoylcarnitine (up) (+) | 201.2, 152.4* | 29.0*, 18.2* | 114.6, 62.6* |
| C18 \| Octadecanoylcarnitine (up) (+) | 4.6, 4.8 | 3.9, 1.2 | 3.4, 1.4 |
| C18:2 \| Linoleoylcarnitine (up) (+) | 15.1, 21.6* | 7.5, 6.0* | 11.8, 13.1* |
| C5 \| Isovalerylcarnitine (up) (+) | 196.2*, 39.4* | 31.8*, 13.6* | 175.1*, 35.7* |
| C5DC \| Glutarylcarnitine (up) (+) | 27.0*, 89.5* | 5.9*, 14.7* | 25.5*, 107.0* |
| C6 \| Hexanoylcarnitine (up) (+) | 107.6*, 137.9* | 27.5*, 43.0* | 80.4*, 99.3* |
| C8 \| Octanoylcarnitine (up) (+) | 89.4*, 100.9* | 24.3*, 24.9* | 90.2*, 80.3* |
| Glutaric acid (up) (+) | 1.9*, 2.1* | 1.2*, 2.0* | 1.5*, 1.5* |
| C18:1 \| Oleoylcarnitine (up) (+) | 8.3, 13.2* | 4.9, 2.9* | 7.0, 4.4 |
| **Homocystinuria N=3** | | | |
| Homocysteine (up) (+) | 0.8, 1.3*, 0.2 | 2.2*, 1.0, 0.3 | 0.8, 1.1, 0.4* |
| Methionine + Methioninesulfoxide (up) (-) | 16.9*, 8.6*, 125.3* | 7.7*, 6.8*, 34.9* | 12.8*, 7.0*, 115.2* |
| Methionine + Methioninesulfoxide (up) (+) | 7.9*, 7.1*, 46.9* | 8.5*, 5.0*, 15.8* | 7.1*, 5.1*, 56.5* |
| **Isovaleric acidemia N=1** | | | |
| C5 \| Isovalerylcarnitine (up) (+) | 62.6* | 33.6* | 41.9* |
| **Long-chain-3-hydroxyacyl CoA dehydrogenase deficiency N=2** | | | |
| Adipic acid (1) (up) (-) | 2.3, 1.7 | 1.0, 0.8 | -0.3, -0.6 |
| Sebacic acid (up) (-) | 6.3, 2.9* | 3.6, 1.2 | 0.3, -0.7* |

| | | | |
|---|---|---|---|
| Sebacic acid (up) (+) | 6.0, 2.9* | 3.6, 1.3 | 0.3, -0.7 |
| Suberic acid (up) (-) | 3.2, 2.3* | 1.2, 0.9 | -0.5, -0.8* |
| Suberic acid (up) (+) | 3.7, 2.7* | 1.4, 1.0 | -0.4, -0.7 |
| **Lysinuric protein intolerance N=2** | | | |
| Arginine (down) (-) | -1.3*, -2.6* | -1.1*, -2.1* | -0.9*, -1.9* |
| Arginine (down) (+) | -1.4*, -1.9* | -1.1*, -1.9* | -1.3*, -1.6* |
| Glutamine + Glutamic acid / N-Methyl-D-Aspartic acid (up) (-) | 3.0*, 3.4* | 0.7*, 6.6* | 0.5, 4.5* |
| Lysine (down) (-) | -0.1, 3.0* | -0.4*, 4.3* | 0.3, 8.0* |
| Lysine (down) (+) | -1.7*, -2.1* | -2.4*, -3.2* | -2.5*, -2.2* |
| Ornithine (down) (-) | 26.6*, 5.2* | 6.2*, 4.4* | 11.7*, 5.2* |
| Ornithine (down) (+) | 0.8, -1.7* | -0.9, -3.1* | -0.3, -2.3* |
| **Malonyl-Coa decarboxylase deficiency N=1** | | | |
| Malonic acid (up) (-) | -0.1 | -0.1 | -0.5 |
| Malonic acid (up) (+) | -0.1 | -0.1 | -0.3 |
| **Maple syrup urine disease N=2** | | | |
| (allo)Isoleucine (up) (-) | 3.3*, 16.9* | 3.1*, 8.7* | 3.6*, 10.3* |
| (allo)Isoleucine (up) (+) | 3.7*, 16.3* | 3.6*, 9.1* | 4.8*, 11.6* |
| 2-Keto-3-methylvaleric acid (up) (-) | -1.6*, 24.7* | -1.8*, 14.7* | -1.0*, 19.4* |
| 2-Keto-4-methylvaleric acid (up) (-) | -2.1*, 2.0 | -3.0*, -0.1 | -1.4*, 0.0 |
| Leucine (up) (-) | 2.3*, 33.4* | 2.6*, 12.2* | 3.4*, 17.7* |
| Leucine (up) (+) | 4.0*, 21.9* | 3.2*, 12.9* | 4.1*, 17.3* |
| Valine (up) (-) | 0.1, 6.7 | 0.8, 3.2* | 1.5, 3.0 |
| **Medium Chain Acyl-CoA Dehydrogenase Deficiency N=4** | | | |
| Adipic acid (1) (up) (-) | 1.4*, 0.6, 0.9, -0.2 | 0.4, -0.1, 0.4, -0.6* | 0.4*, 0.1, 0.3, -0.4 |
| C10:1 \| Decenoylcarnitine (up) (+) | 43.6*, 33.3*, 8.8*, 26.5* | 21.6*, 18.1*, 7.4*, 15.6* | 56.2*, 44.6*, 12.1*, 35.1* |
| C6 \| Hexanoylcarnitine (up) (+) | 52.2*, 45.2*, 11.1*, 31.5* | 19.8*, 18.1*, 6.8*, 14.0* | 69.5*, 59.4*, 14.7*, 41.1* |
| C8 \| Octanoylcarnitine (up) (+) | 51.5*, 65.5*, 27.9*, 36.0* | 23.9*, 26.7*, 15.9*, 18.4* | 110.2*, 134.2*, 57.2*, 73.0* |
| Hexanoic acid / Trans-cyclohexane-1,2-diol (up) (-) | 0.5, 0.3, 0.6, 0.7 | -0.3, -1.2*, 0.4, 1.0 | -0.1, -0.2, 0.0, 0.1 |
| Sebacic acid (up) (-) | 0.0, 0.6, 0.4, -0.6 | -0.5, -0.2, -0.1, -0.6 | -0.3*, 0.1, -0.0, -0.5 |
| Sebacic acid (up) (+) | 0.3, 0.6, 0.7, -0.5 | -0.3, -0.2, 0.1, -0.5 | 0.3*, 0.5, 0.5, -0.0 |
| Suberic acid (up) (-) | 0.9*, 0.9, 0.8*, -0.3 | 0.0, -0.0, 0.2, -0.6 | 0.3*, 0.4, 0.3, -0.3 |
| Suberic acid (up) (+) | 0.9, 1.0, 0.5, -0.2 | 0.2, -0.1, -0.0, -0.5 | 0.5, 0.5, 0.3, -0.1 |
| **Methylmalonyl-CoA mutase deficiency N=1** | | | |
| C3 \| Propionylcarnitine (up) (+) | 111.6* | 34.1* | 181.0* |

| | | | |
|---|---|---|---|
| **Organic cation transporter 2 deficiency N=1** | | | |
| C0 \| L-Carnitine (down) (-) | -1.9* | -0.7 | -1.2* |
| C0 \| L-Carnitine (down) (+) | -2.3* | -1.4* | -0.9 |
| **Ornithine aminotransferase N=1** | | | |
| Guanidinoacetic acid (down) (+) | -1.4* | -2.5* | -1.6* |
| Ornithine (up) (-) | -0.3 | -0.5* | -0.6 |
| Ornithine (up) (+) | 42.1* | 11.8* | 37.9* |
| **Ornithine transcarbamylase deficiency N=2** | | | |
| Citrulline (down) (-) | 3.4*, 1.4* | 1.2*, 0.5 | 1.9*, 1.0* |
| Citrulline (down) (+) | 1.8*, 2.4* | 0.5*, 0.9 | 1.5*, 1.4* |
| Glutamine + Glutamic acid / N-Methyl-D-Aspartic acid (up) (-) | 1.7*, 1.0* | 0.8*, 0.9 | 0.4, 1.7* |
| Uridine (up) (-) | 7.3*, 5.1* | 4.1*, 4.5* | 5.9*, 4.9* |
| **Phenylketonuria N=3** | | | |
| N-Acetylphenylalanine (up) (-) | 127.6*, 41.1, 124.6* | 24.5*, 11.7*, 32.7* | 40.4*, 13.1, 55.5* |
| Phenylalanine (up) (-) | 88.9*, 47.5*, 163.7* | 33.6*, 23.1*, 43.2* | 86.0*, 46.9*, 119.7* |
| Phenylalanine (up) (+) | 49.6*, 33.3*, 76.8* | 20.8*, 16.4*, 29.6* | 53.7*, 37.1*, 67.0* |
| Phenyllactic acid (up) (-) | 126.2*, 77.1*, 177.3* | 63.1*, 48.1*, 10.1* | 407.0*, 249.5*, 19.2* |
| alpha-N-Phenylacetylglutamine (up) (-) | 2.4*, 1.6*, 5.7* | 2.1*, 1.5*, 2.7* | 3.1*, 2.0, 3.7* |
| alpha-N-Phenylacetylglutamine (up) (+) | 3.1*, 2.0, 4.7* | 2.8*, 1.9*, 2.7* | 4.0*, 2.4, 3.2* |
| **Propionic acidemia N=1** | | | |
| C3 \| Propionylcarnitine (up) (+) | 113.6* | 41.2* | 115.6* |
| **Thymidine phosphorylase deficiency N=1** | | | |
| Thymidine (up) (-) | 57.7* | 104.7* | 47.3* |
| **Tyrosinemia I N=2** | | | |
| 4-Hydroxyphenyllactic acid (up) (-) | 337.5*, 1047.6* | 11.8*, 16.5* | 360.7*, 567.0* |
| 4-Hydroxyphenylpyruvic acid (up) (-) | 2.8*, 1.4 | 1.9*, 1.4 | 3.0*, 0.3 |
| N-Acetyltyrosine (up) (-) | 28.2*, 89.9* | 19.7*, 32.3* | 60.8*, 43.4* |
| Tyrosine (up) (+) | 16.6*, 40.2* | 8.6*, 20.9* | 22.4*, 37.9* |
| **Very Long Chain Acyl-CoA Dehydrogenase Deficiency N=1** | | | |
| C14 \| Tetradecanoylcarnitine (up) (+) | 47.9* | 11.1* | 39.0* |
| C14:1 \| Tetradecenoylcarnitine (up) (+) | 81.2* | 23.4* | 165.2* |
| C16 \| Hexadecanoylcarnitine (up) (+) | 6.7* | 2.0* | 2.5* |
| C18 \| Octadecanoylcarnitine (up) (+) | 3.1* | 1.1* | 1.0* |
| c18:1 \| Oleoylcarnitine (up) (+) | 5.7* | 2.0* | 2.7* |

| | | | |
|---|---|---|---|
| **Carnitine palmitoyltransferase II N=2** | | | |
| Adipic acid (1) (up) (-) | 0.5, 0.3 | 0.2, 0.1 | 0.6, 0.3 |
| C0 \| L-Carnitine (down) (-) | 0.4, -0.6 | -0.1, -1.1 | 1.2*, -0.7 |
| C0 \| L-Carnitine (down) (+) | 0.9, -0.3 | 1.3, -0.8* | 0.7, -0.4 |
| C16 \| Hexadecanoylcarnitine (up) (+) | 0.5, 19.7* | 2.0*, 5.8* | 2.0*, 11.8* |
| C18 \| Octadecanoylcarnitine (up) (+) | 1.7*, 29.8* | 3.7*, 7.5* | 4.0*, 20.0* |
| Sebacic acid (up) (-) | 0.4, 0.1 | 0.1, -0.2 | 0.4, -0.1 |
| Sebacic acid (up) (+) | -0.9*, 0.3 | 0.4, -0.1 | 0.5*, -0.1 |
| Suberic acid (up) (-) | 0.8, 0.2 | 0.5, -0.2 | 0.7, -0.0 |
| Suberic acid (up) (+) | -0.7, 0.0 | 0.6, -0.3 | 0.6, -0.1 |
| C18:1 \| Oleoylcarnitine (up) (+) | -0.1, 20.0* | 1.6, 4.9* | 2.5, 12.3 |

### S6 Performance *BC-Metchalizer* for a different number of internal standards

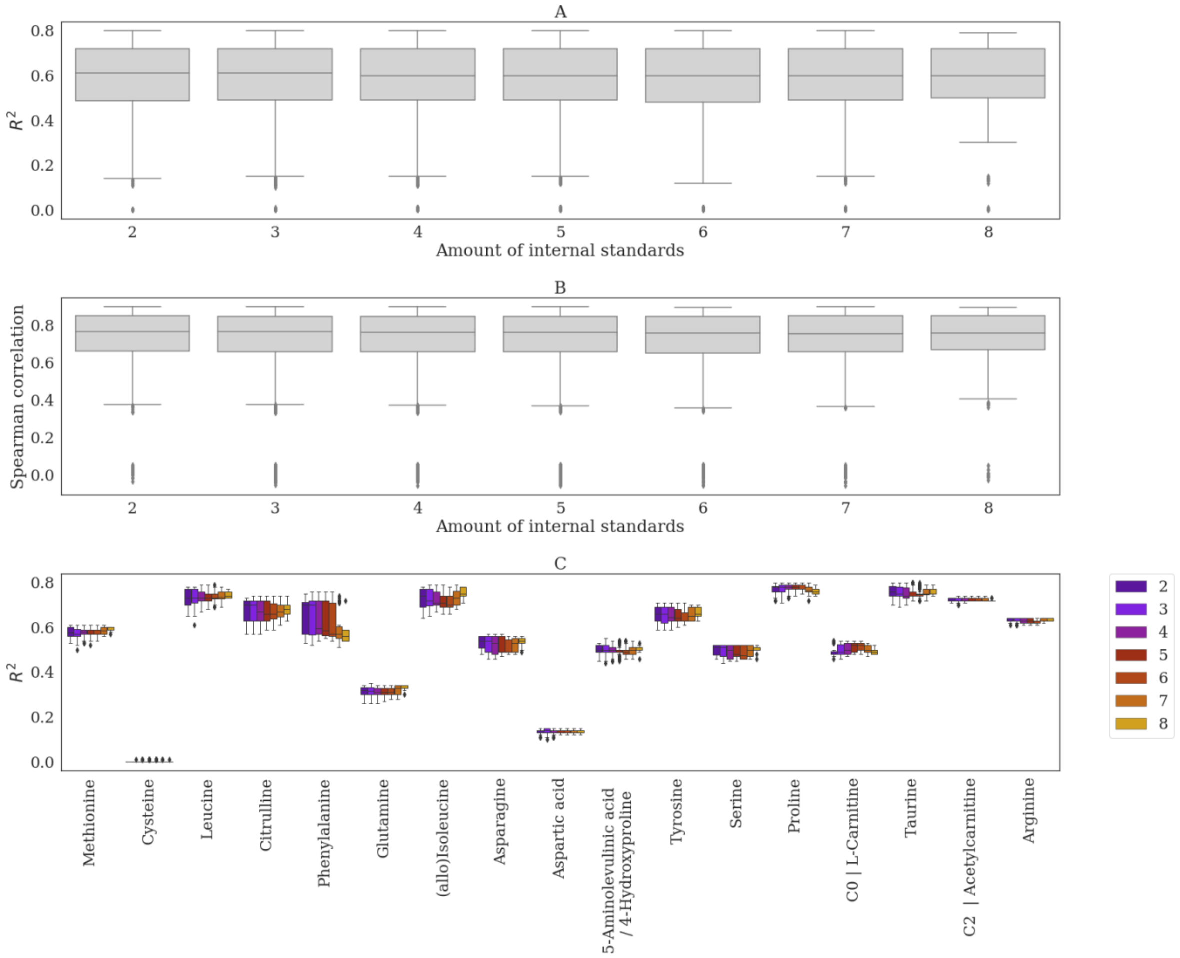

Figure 7. Performance of normalization based on a comparison with quantitative measurements for increasing the number of internal standards with *BC-Metchalizer*. A) $R^2$ B) Spearman correlation C) $R^2$ per metabolite and number of internal standards. The legend indicates how many internal standards were used for normalization.

Since *BC-Metchalizer* uses multiple internal standards we explored the influence of the number of internal standards on normalization by comparing quantitative measurements with the abundancies after normalization. Subfigure in A, B in Figure 5 shows the overall Coefficient of determination $R^2$, Spearman correlation, for the quantified metabolites (see subfigure C). Clearly, no increase/decrease in performance was observed when increasing the number of internal standards. However, looking at the individual normalization performances per metabolite showed that some combinations of internal standards improved or decreased $R^2$. For example, phenylalanine showed most variation in performance and more internal standards did not lead to increased performance. From here, it seemed that some combinations of internal standards normalized certain metabolites better.

### S7 Performance *BC-Metchalizer* for a different amount of batches

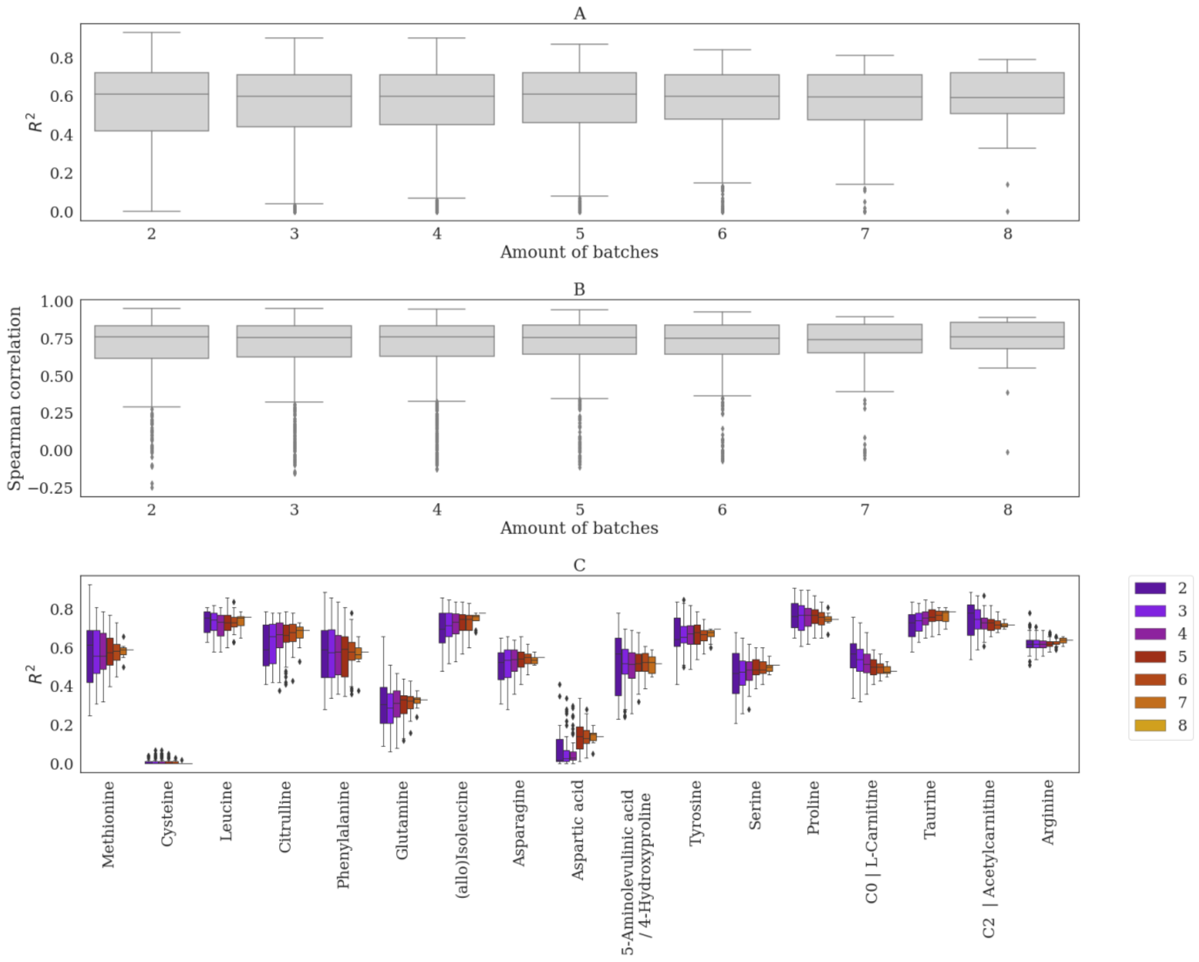

Figure 8. Performance of normalization based on a comparison with quantitative measurements for increasing the amount of batches with *BC-Metchalizer*. A) $R^2$ B) Spearman correlation C) $R^2$ per metabolite and amount of batches. The legend indicates how many batches were used for normalization. 9 internal standards were used for normalization.

Robustness of normalizing with BC-*Metchalizer* was investigated by normalizing a different number of batches (and all of their combinations). Ideally, the performance of normalization should be constant over the amount of batches being analyzed. Subfigure in A, B in Figure 6 shows the overall Coefficient of determination $R^2$, Spearman correlation, for the quantified metabolites (see subfigure C) for increasing batches. It can be observed that globally its performance was constant over the whole range. However, on the metabolite specific level we do observe difference between the amount of batches being used.

### S8 Biomarker detection performance for age-dependent biomarkers

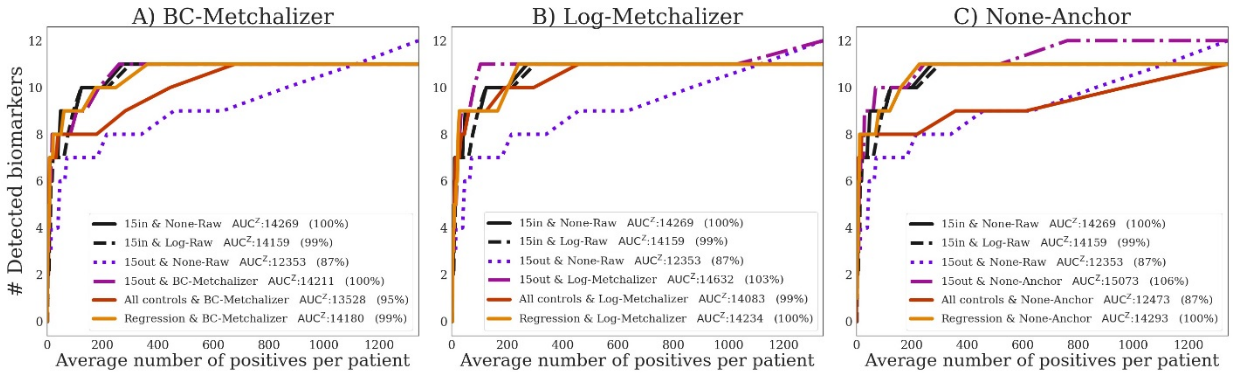

Figure 9. The number of detected <u>age-dependent</u> biomarkers (see S3 Table 3) versus the average number of positives per patient. The average number of positives was determined from all features i.e. 758 positively - and 583 negatively ionized feature (excluding standards). A curve in each (sub)figure was formed by increasing $Z_{abnormal}$ (Methods). The legend indicates (per curve) the methods used to determine Z-scores and how data was normalized, the AUC and AUC expressed as percentage of the AUC of *15in&None-Raw*. Performances using A) *BC-Metchalizer*, B) *Log-Metchalizer*, C) *None-Anchor*.

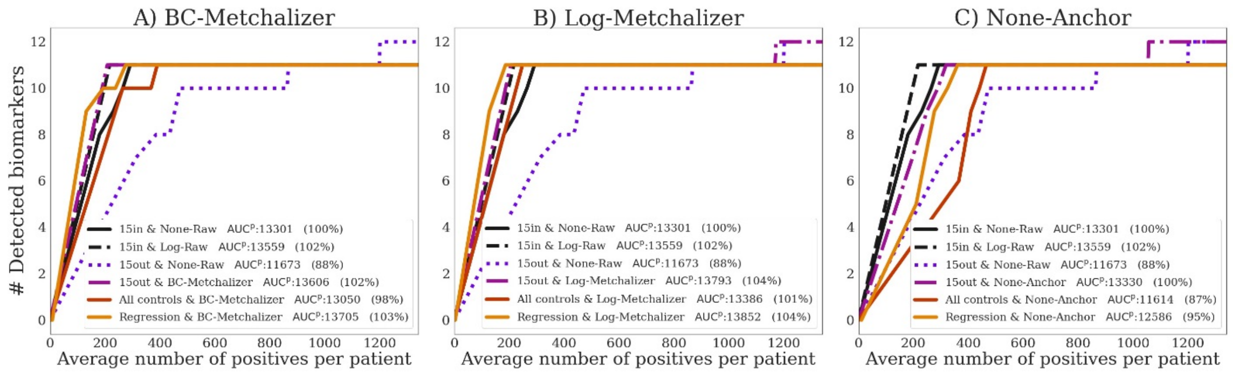

Figure 10. The number of detected <u>age-dependent</u> biomarkers versus the average number of positives per patient. A curve in each (sub)figure was formed by increasing the p-values (Methods). The legend indicates (per curve) the methods used to determine Z-scores and p-values and how data was normalized, the AUC and AUC expressed as percentage of the AUC of *15in&None-Raw*. Performances using A) *BC-Metchalizer*, B) *Log-Metchalizer*, C) *None-Anchor*.

Figure 9 and Figure 10 show the biomarker detection performance for different normalization method and approaches to calculate Z-scores ( as indicated by the legends). Here, we included only age-dependent biomarkers: Guanidinoacetic acid(+), Homoarginine(-) and N-Acetyltyrosine(-) , 2-Ketoglutaric acid(-), Citrulline(-) and Ornithine(-). We observe that using *Regression* seemed to outperform *All controls*.

### S9 Biomarker detection performance for 8 normalization methods using Z-scores

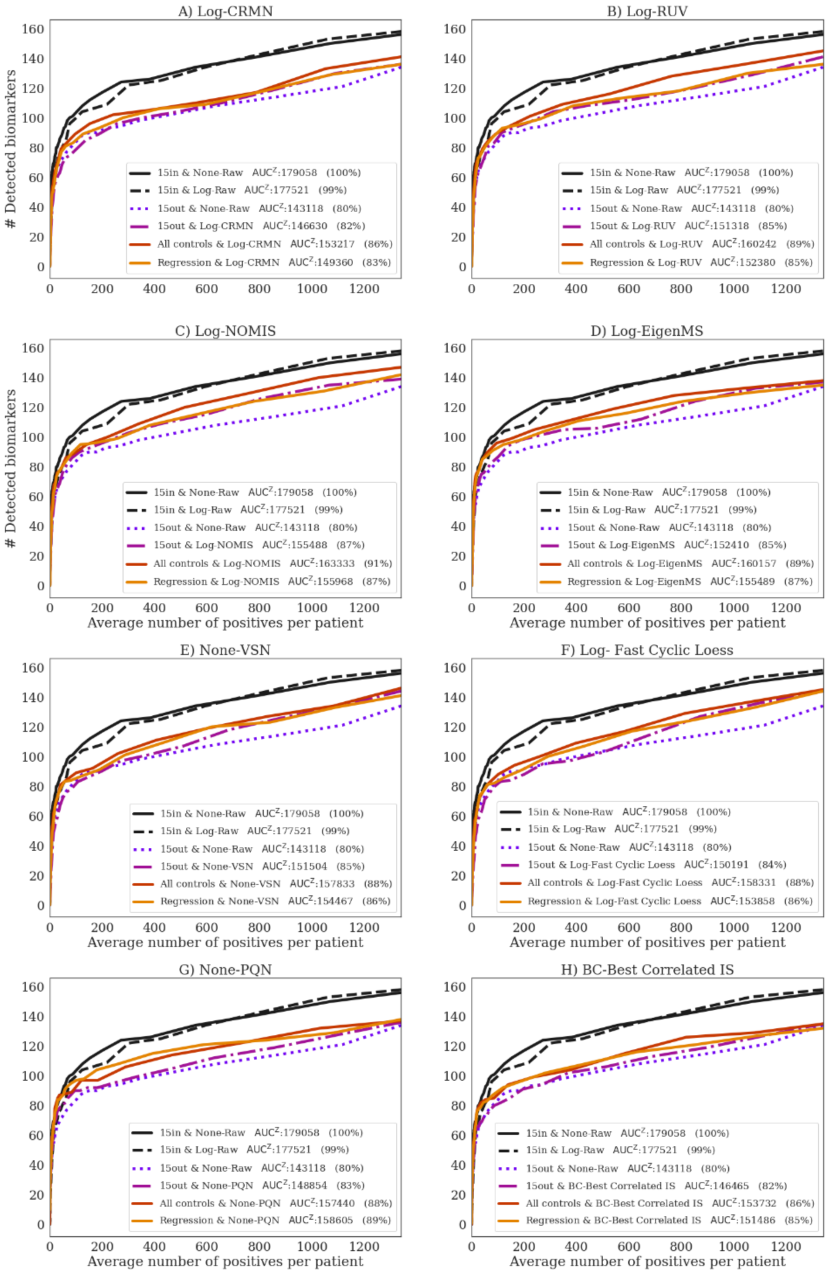

Figure 11. The number of detected biomarkers versus the average number of positives per patient using the Z-scores. A) Performances using *Log-CRMN*. B) Performances using *Log-RUV*. C) Performances using *Log-NOMIS*. D) Performances using *Log-EigenMS*. E) Performances using *None-VSN*. F) Performances using *Log- Fast Cyclic Loess*. G) Performances using *None-PQN*. H) Performances using *BC-Best Correlated IS*.

### S10 The influence of log-transformation on Z-scores and p-values

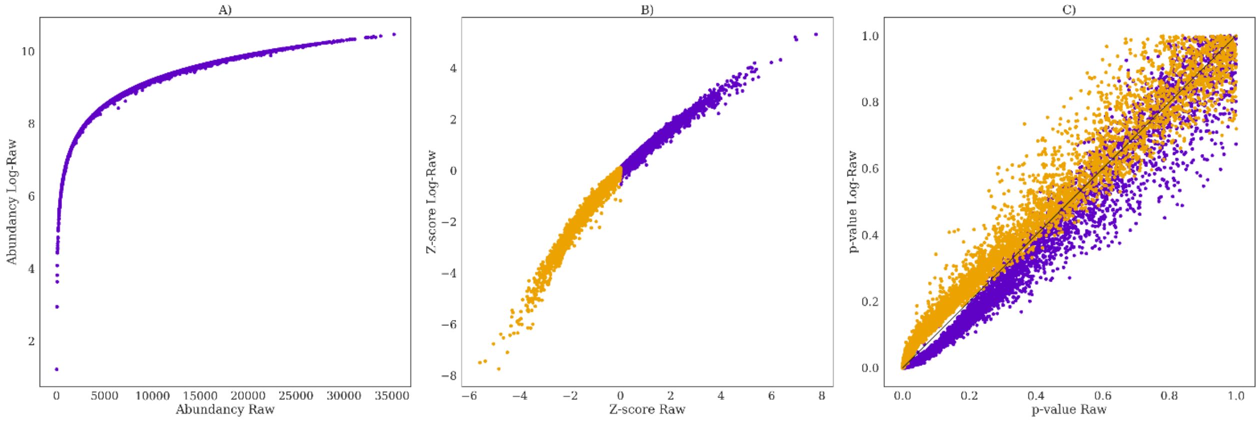

Figure 12. A) Shows simulated log-transformed abundancies (*Log-Raw*) versus the raw abundancies (*Raw*). B) The difference in Z-scores between *Log-Raw* and *None-Raw*. C) The difference in Welch's t-test p-values between *Log-Raw* and *None-Raw*. Simulations were performed by randomly drawing a mean $\mu_{\text{pop}}$ = Normal(mu = 10000, sigma = 5000) followed by 15 random 'controls' samples from Normal(mu = $\mu_{\text{pop}}$, sigma = $\mu_{\text{pop}}$ / 5). From the same normal distribution we drew $\mu_{\text{trip}}$, and three 'triplicate measurements' were drawn from Normal(mu = $\mu_{\text{trip}}$, sigma = $\mu_{\text{trip}}$ / 5). Z-scores were calculated from the average en standard deviation obtained from the 15 'controls'.
